# Fast and accurate construction of multiple sequence alignments from protein language embeddings

**DOI:** 10.64898/2026.01.02.697423

**Authors:** Minh Hoang, Isabel Armour-Garb, Mona Singh

## Abstract

Multiple sequence alignment (MSA) is a foundational task in computational biology, under-pinning protein structure prediction, evolutionary analysis, and domain annotation. Traditional MSA algorithms rely on pairwise amino acid substitution matrices derived from conserved protein families. While effective for aligning closely related sequences, these scoring schemes struggle in the low-identity “twilight zone.” Here, we present a new approach for constructing MSAs leveraging amino acid embeddings generated by protein language models (PLMs), which capture rich evolutionary and contextual information from massive and diverse sequence datasets. We introduce a windowed reciprocal-weighted embedding similarity metric that is surprisingly effective in identifying corresponding amino acids across sequences. Building on this metric, we develop ARIES (**A**lignment via **R**ec**I**procal **E**mbedding **S**imilarity), an algorithm that constructs a PLM-generated template embedding and aligns each sequence to this template via dynamic time warping in order to build a global MSA. Across diverse benchmark datasets, ARIES achieves higher accuracies than existing state-of-the-art approaches, especially in low-identity regimes where traditional methods degrade, while scaling almost linearly with the number of sequences to be aligned. Together, these results provide the first large-scale demonstration of the power of PLMs for accurate and scalable MSA construction across protein families of varying sizes and levels of similarity, highlighting the potential of PLMs to transform comparative sequence analysis.

## Introduction

Comparative analysis of protein sequences provides fundamental insights into their structure, function, and evolution. A central tool for such analysis is multiple sequence alignment (MSA), which arranges evolutionarily-related residues across homologous sequences to reveal conserved positions and motifs. The quality of an MSA directly influences a wide range of downstream applications, including protein structure prediction (Marks et al. 2012), phylogenetic reconstruction (Whelan and Goldman 2001), and functional annotation (Capra and Singh 2007). In modern structure prediction systems such as AlphaFold (Jumper et al. 2021), co-variation patterns extracted from MSAs provide essential constraints for learning residue–residue contacts and 3D geometry, making accurate alignments a prerequisite for many advances in computational proteomics.

Over the years, numerous MSA algorithms have been developed (e.g., Thompson et al. (1994); Katoh et al. (2002); Do et al. (2005); Sievers et al. (2011); Liu et al. (2012); Mirarab et al. (2015)). Most of these approaches follow a progressive alignment strategy where the most similar sequences are aligned first, and more distant sequences are iteratively added according to a guide tree. Alignment scores are typically computed using substitution matrices such as PAM (Dayhoff et al. 1978) and BLOSUM (Henikoff and Henikoff 1992), which are comprised of log-odds scores reflecting how often one residue substitutes for another (Altschul 1991), as estimated from conserved protein families. MSAs built using these matrices are highly accurate when sequence identity is high, but lose reliability in the low sequence identity “twilight zone” of alignment (Rost 1999). A major limitation stems from the context-independent nature of traditional substitution matrices, which assign each residue the same fixed similarity score for a given substitution, regardless of its biochemical or structural environment within the sequence.

The advent of protein language models (PLMs) (Bepler and Berger 2021; Alley et al. 2019; Rao et al. 2019; Bepler and Berger 2019; Elnaggar et al. 2021; Rives et al. 2021) offers a transformative opportunity to overcome this limitation. PLMs such as ESM-2 (Lin et al. 2023) and ProtT5 (Elnaggar et al. 2021) are trained on massive protein sequence databases to learn context-aware embeddings that implicitly capture underlying evolutionary relationships. PLM embeddings have been used for BLAST-like database searches, where homologs are identified based on sequence-level embedding similarity (Schütze et al. 2022; Kilinc et al. 2023; Kaminski et al. 2023; Iovino et al. 2024). In recent years, several studies have extended the use of PLM embeddings to construct MSAs, with varying degrees of success.

The first such approach, vcMSA (McWhite et al. 2023), employs a clustering-based strategy to recursively assemble aligned columns from embedding representations, but its accuracy and scalability remain limited for large sequence sets. Meanwhile, methods such as EBA (Pantolini et al. 2024) and PEbA (Iovino and Ye 2024) incorporate embedding-based similarity measures directly into classical pairwise alignment dynamic programming frameworks, but they lack mechanisms to reconstruct full MSAs. More recently, learnMSA2 (Becker and Stanke 2024) integrates PLM embeddings into a hidden Markov model-based framework for aligning large, well-conserved families. However, its statistical estimates become unstable for small or highly divergent sets, leading to substantial drops in accuracy. Collectively, these PLM-based methods have shown that sequences can be aligned via PLM-derived substitution scores, but none have yet consistently and efficiently yielded high-quality alignments across both small and large protein families. We provide brief descriptions of these methods in Appendix B.

In this paper, we present a robust and scalable PLM-based technique for constructing MSAs and demonstrate its excellent performance across diverse benchmarks of varying sizes and degrees of heterogeneity. Our approach introduces three key innovations: (1) a context-aware, reciprocal similarity measure that enhances the robustness of residue comparisons; (2) a scalable two-phase strategy, based on “star” alignment (Altschul and Lipman 1989), that builds MSAs from pairwise alignments with a template sequence without requiring explicit gap penalty functions; and (3) a template-construction procedure that leverages PLMs to generate a well-informed central sequence for this star alignment strategy.

Our similarity measure is motivated by the observation that prior residue-level PLM-derived similarities (McWhite et al. 2023; Iovino and Ye 2024) can struggle to identify aligned residue pairs within low-identity regions of the sequences. To reduce sensitivity to local context perturbations, we incorporate a windowing step that aggregates similarity over the respective local windows of the residue pair being compared. To further distinguish true evolutionary correspondences from superficial contextual matches, we introduce a reciprocal weighting mechanism that prioritizes mutually recognized residue pairs over asymmetric matches (see **Methods**). This reciprocality reward sharpens the alignment signal and proves effective in the low-identity regime (see **Ablation studies**).

Using our similarity measure, we leverage dynamic time warping (DTW) as the core alignment primitive in our workflow (Sakoe and Chiba 1978). DTW offers a key advantage over traditional alignment algorithms: it naturally handles insertions and deletions without requiring explicit gap penalties, instead representing them as many-to-one or one-to-many mappings along the alignment path. This feature is especially useful in the context of PLM-based alignment, since gap embeddings cannot be generated without knowing their locations in advance. Building on this primitive, we construct MSAs of protein families based on a star alignment framework (Fig. 1, top), where each protein is pairwise-aligned against a representative template sequence. The resulting pairwise alignments are merged into a global MSA by grouping residues that map to the same template position, with many-to-one cases disambiguated by selecting the correspondences with highest embedding similarities.

**Fig. 1.**
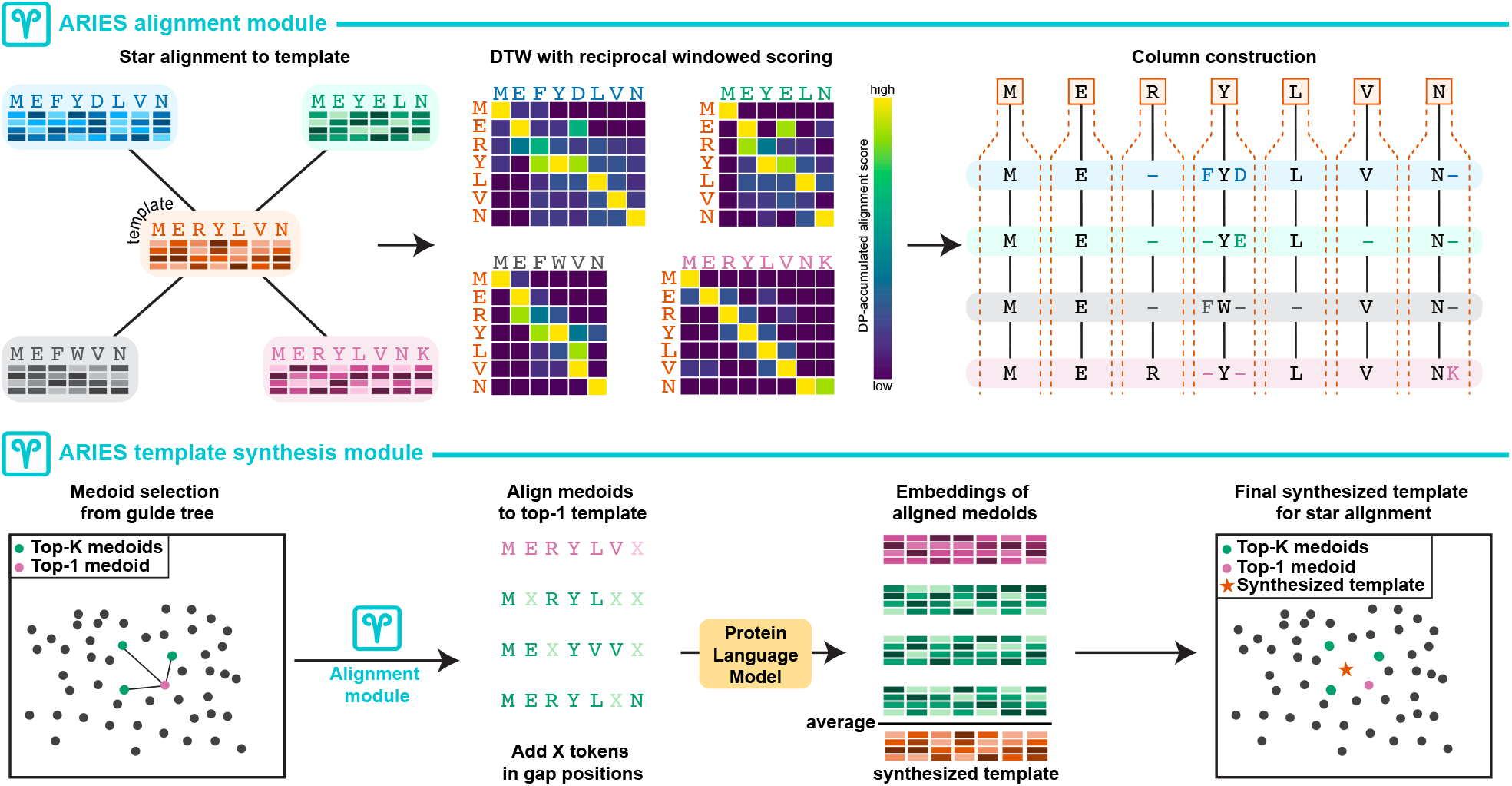
The algorithmic core of ARIES consists of two modules, the *star-alignment* module (top) and the *template synthesis* module (bottom). In the alignment module, all sequences are aligned to a single template via dynamic time warping (DTW) using a reciprocal, window-based embedding similarity score. Per-template-position column blocks are constructed by selecting anchor residues (denoted by vertical black lines), placing remaining residues around these anchors, and adding gaps to form coherent alignment columns. In the template synthesis module, the *K* sequences most similar to all other sequences (i.e., the top-*K* medoids) are aligned using the ARIES alignment module with the top-1 medoid as the template. Gaps in the alignment are replaced with ‘X’ tokens, and the sequences with ‘X’ tokens are individually re-embedded with the PLM. A synthesized consensus template is produced by position-wise averaging of the resulting embeddings across all *K* sequences, including the top-1 medoid.

For extremely large sequence sets, however, multiple subgroups (or clusters) of highly similar sequences are often observed, and simply selecting a single sequence from the input set favors certain subgroups, and hence degrades overall performance (see **Ablation studies**). To overcome this shortcoming of classical star alignment (Altschul and Lipman 1989), we employ PLMs to synthesize a balanced and informative template that incorporates features from the top-*K* “medoid” sequences. Specifically, in the first phase of our alignment approach, we identify the *K* sequences that are the closest to all other sequences, and then perform a miniature star alignment, using the global medoid as an initial template and obtaining a coarse alignment for these *K* sequences (Fig. 1, bottom). In the second phase, we re-embed the aligned medoid sequences individually with a PLM and compute the position-wise average of their embeddings to generate a synthesized template that ideally captures shared evolutionary signals across subgroups. This refined template is subsequently used to align all sequences in the input set.

We refer to this framework as ARIES (**A**lignment via **R**ec**I**procal **E**mbedding **S**imilarity). To assess the practical utility of this framework, particularly in the twilight zone where substitution-matrix-based methods lose reliability, we benchmark ARIES against leading MSA tools across datasets spanning a range of sequence identities and family sizes, measuring both alignment accuracy and computational scaling (see **Results**). These analyses are designed to determine whether PLM-based alignment can extend reliable MSA construction into low-identity regimes while remaining computationally practical for large protein families, an advance that could benefit a wide range of applications, including structure prediction, phylogenetic reconstruction, and functional annotation.

## Methods

### Overview

ARIES builds on the principle of embedding-based pairwise alignment. Amino acid embeddings are obtained for each sequence (see **Embedding generation**), and pairwise correspondences between residues are computed using their contextual similarity in the learned embedding space of PLMs (see **ARIES similarity score matrix**). Pairs of sequences are aligned using the DTW algorithm (see **Dynamic Programming for Pairwise Sequence Alignment**). To scale this pairwise procedure to multiple sequences, we adopt a two-phase star alignment MSA building approach (see **Constructing MSAs from pairwise alignments with template sequence**, Fig. 1), which first generates a template sequence (see **Constructing the template sequence**), and then aligns each remaining sequence to this template using DTW. From these pairwise alignments, we reconstruct the global MSA by iteratively building column blocks centered around each template residue (see **Star alignment procedure**). As DTW can produce many-to-one mappings, we further introduce a disambiguation step to infer residue correspondences and consistent gap placements (see **Gap inference**).

### Embedding generation

We obtain embeddings of protein sequences using PLMs such as ESM-2 (Lin et al. 2023) and ProtT5 (Elnaggar et al. 2021). Given an amino acid sequence (*a*_1_, *a*_2_, …, *a*_*L*_), the PLM produces a set of hidden states

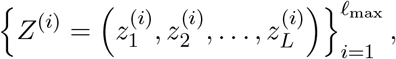

where *l*_max_ is the number of layers in the PLM and 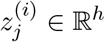 denotes the *h*-dimensional embedding of *a*_*j*_ at the layer *i*. By convention, *Z*^(1)^ denotes the embedding representation at the final layer (prior to the output logit). Special tokens added by the PLM (e.g., CLS or EOS) are not considered in our framework. Since different layers of PLMs capture complementary structural and functional information (Lupo et al. 2022), we represent each residue *a*_*j*_ as the concatenation of its hidden embeddings from the last *l* layers (*l* ≤ *l*_max_): 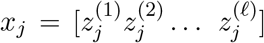, resulting in a vector with *d* ≜ *lh* dimensions. We give the details of PLMs used in our experiments in Appendix C, and ablate the effect of varying *l* in a later section (see **Ablation studies**).

### ARIES similarity score matrix

Unlike traditional substitution matrices such as BLOSUM (Henikoff and Henikoff 1992) or PAM (Dayhoff et al. 1978), PLM embeddings capture dependencies between residues through self-attention pre-training, which enables them to reflect subtle contextual information such as residue co-evolution, local structure, and domain-specific motifs (Lupo et al. 2022). Deriving similarity scores directly from these embeddings thus allows the alignment process to operate in a semantically meaningful representation space, where geometric proximity reflects functional or structural correspondence.

Let 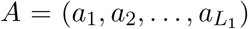 and 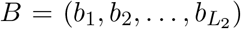 be two protein sequences with their respective PLM embeddings *X* = (*x*_1_, *x*_2_, …, *x*_*L*_ ) 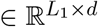 and 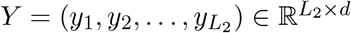. Their residue-level similarity can be encoded as a matrix 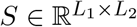, where each entry *S*_*ij*_ quantifies the similarity between the pair of amino acids *a*_*i*_ and *b*_*j*_. We can measure this similarity using the negative Euclidean distance (NED) of their embeddings, i.e., *S*_*ij*_ ≜ −∥*x*_*i*_ − *y*_*j*_∥_2_. Pantolini et al. (2024) demonstrated that Euclidean distance over residue embeddings can effectively capture similarity for use in a downstream alignment algorithm. While the NED metric can by itself, with no alignment step, help identify amino acid pairs across sequences that should be aligned (see **Reciprocal, window-based embedding similarity enables detection of aligned residues**), to obtain a more stable and discriminative similarity function, we extend NED in two ways: first by aggregating information over local windows of amino acids, and second by applying a reciprocal weighting scheme that prioritizes mutually consistent matches. These augmentations are described in what follows.

#### Window-based similarity measure

Since the NED metric can sometimes lead to high similarities between amino acid pairs that should not be aligned (see Fig. 2), we first generalize NED to capture similarity between windows of embeddings. This approach is motivated by prior work showing that local context can improve both PLM-based representations and sequence alignment. We previously demonstrated that locality-aware pooling of PLM embeddings improves downstream prediction (Hoang and Singh 2025), while CS-BLAST (Biegert and Söding 2009) showed that context-specific similarity scores derived from local sequence windows improve alignment quality over context-independent substitution scores. Here, we adapt this principle of incorporating local context to construct a local-context, residue-pairwise similarity score for MSA. To the best of our knowledge, no prior embedding-based MSA method uses a windowed similarity measure of this form. For some choice of window span *w*, we encode each residue *a*_*i*_ in *A* by concatenating all embeddings in the (2*w* + 1)-length context window centered around it, i.e., 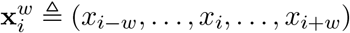. Likewise, each residue *b*_*j*_ in *B* is encoded by 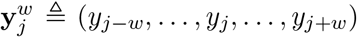. To ensure that each residue is associated with a sufficiently large context window, we pad both ends of the original sequence with *w* contiguous “unknown amino acid” tokens, represented by the token ‘X’ in most PLM vocabularies. The window-level similarity matrix between *A* and *B* is then given by the matrix 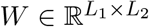, where each entry *W*_*ij*_ is obtained as a weighted aggregation over corresponding token-pair similarities between 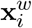 and 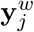. Specifically, we define:

**Fig. 2.**
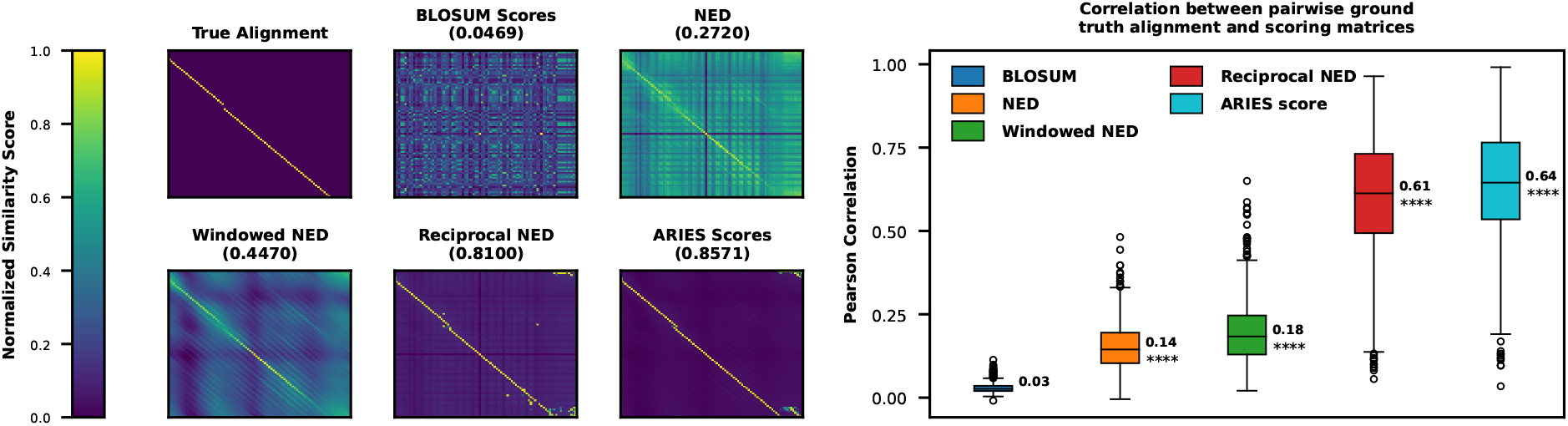
*Left:* Binary true alignment matrix of two BAliBASE RV11 sequences, followed by heatmaps of five different pairwise residue-similarity matrices for these sequences with Pearson correlation of these similarity scores with the true alignment matrix given. *Right:* Box plots of Pearson correlations between the true alignment and the same five scoring schemes across all sequence pairs in BAliBASE RV11. Median correlation is annotated next to each box. Stars denote statistically significant higher values over than the method to the left (*p ≤* 1*e*^*−*4^, one-sided Wilcoxon signed-rank test with Holm-Bonferroni correction).

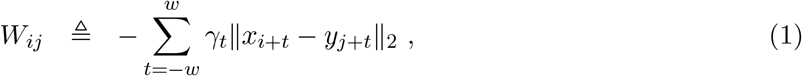

where 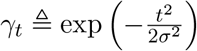 follows a discrete, zero-centered Gaussian kernel, assigning greatest importance to the central residue and progressively discounting the contributions of neighboring positions.

#### Reciprocal-weighted similarity measure

While window averaging reduces noise, contextual differences across the embedding space of *X* and *Y* can give rise to bias in their pairwise similarity scores. We find that some residues produce broad, non-specific similarity patterns, showing moderate similarity to many positions in the other sequence. Conversely, other residues align strongly to only a few positions (see Fig. 2). Such asymmetry can mislead the alignment process, as residues with non-specific similarities may dominate the scoring matrix and cause incorrect matches (see **Reciprocal, window-based embedding similarity enables detection of aligned residues**). To counteract this phenomenon, we further introduce a reciprocal-weighted similarity term that adjusts entries in *W* to reward mutually consistent correspondences.

The mutual consistency of two residues *a*_*i*_ and *b*_*j*_ can be assessed by weighing their alignment signal *W*_*ij*_ against other entries of *W* in the same row and column. Specifically, we compute the softmax normalizations of *W*_*ij*_ along row *i* and column *j* of *W* :

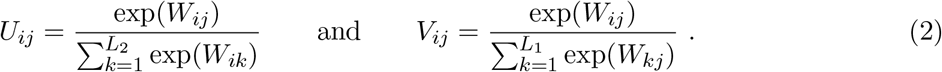

Here, *U*_*ij*_ represents how strongly *a*_*i*_ favors *b*_*j*_ over other residues in *B*, and likewise *V*_*ij*_ reflects the reciprocal preference in the opposite direction. If both perspectives agree, i.e., each residue strongly favors the other compared to all alternatives, then the alignment is considered mutually consistent. Building on this intuition, we define the *reciprocal consistency score* as *R*_*ij*_ ≜ 0.5 × log (*U*_*ij*_*V*_*ij*_), which corresponds to the log of the geometric mean between the two directional affinities. We note that the log transformation guarantees that *R*_*ij*_ is negative, with *R*_*ij*_ becoming more negative with decreasing specificity. Since *W*_*ij*_ is also negative and becomes more so when amino acids are less similar to each other in embedding space, the *W* and *R* matrices can naturally be combined via addition. Our proposed pairwise similarity matrix is thus defined as *S* = *W* + *λR*, where the hyperparameter *λ* controls the strength of reciprocal reinforcement. We investigate the effect of the *λ* hyperparameter in a later section (see **Ablation studies**).

### Dynamic Programming for Pairwise Sequence Alignment

Dynamic programming (DP) forms the computational backbone of most sequence alignment algorithms, with the Needleman-Wunsch algorithm (Needleman and Wunsch 1970) finding the best global alignment of two sequences given a substitution matrix and gap penalty score. Most traditional MSA approaches use heuristic approaches to assign gap penalty scores, though some learn position-specific gap penalties directly from sequence data (e.g., the HHalign method (Remmert et al. 2012) used by Clustal Omega (Sievers et al. 2011)). In the embedding-based alignment paradigm, PLM embeddings directly provide residue–residue similarity scores, but do not provide a corresponding model for insertion and deletion events, making it unclear how gap penalties should be specified. Indeed, existing embedding-based alignment approaches either avoid explicit pairwise alignment altogether (e.g., vcMSA (McWhite et al. 2023) and learnMSA2 (Becker and Stanke 2024)) or depend on ad-hoc gap penalty schemes (e.g., PeBA (Iovino and Ye 2024)). To address this challenge, we adopt the DTW formulation (Sakoe and Chiba 1978) that is frequently used in signal processing domains to align time series data. The DTW framework aligns two time series by allowing local stretching and compression along the time axis to maximize the overall correspondence between aligned time points. When applied to biological sequence alignment, this relaxes the strict one-to-one or one-to-gap constraint of classical alignment methods, yielding gap-free alignment maps in which a single residue may correspond to multiple contiguous residues in the other sequence. In this work, we show that gap placement can subsequently be inferred through a post-hoc refinement procedure. The DP recursion underlying DTW is described below.

#### Definition 1

**(Dynamic Time Warping on Embedding Space)**

*Let* 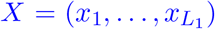 *and* 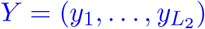 *be two embedding sequences of lengths L*_1_ *and L*_2_, *and* 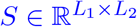 *their pairwise similarity matrix defined previously. A warping path P between X and Y is defined as a sequence of index pairs* (*i, j*) *that starts at* (1, 1), *ends at* (*L*_1_, *L*_2_), *and at each step advances i, j, or both by one. Let* P_*XY*_ *be the set of all such warping paths. The DTW algorithm finds the optimal path* 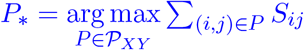 *with maximum total similarity score according to S. Letting D*(*i, j*) *denote the maximum accumulated similarity of any warping path ending at* (*i, j*), *the DP recursion to compute D*(*i, j*) *is given by:*

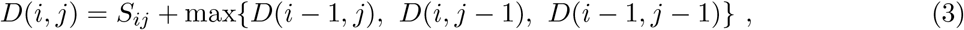

*where we initialize D*(0, 0) = 0 *and D*(*i*, 0) = *D*(0, *j*) = −∞ *for all i* ∈ [*L*_1_] *and j* ∈ [*L*_2_]. *The final warping path, corresponding to the maximum total similarity between the sequences, is recovered by backtracking from* (*L*_1_, *L*_2_) *to* (1, 1), *analogous to traditional sequence alignment (Sakoe and Chiba 1978)*.

We note that DTW is fundamentally different from Needleman-Wunsch (NW) with a 0 gap penalty. NW adds the residue-residue similarity score only for diagonal moves, corresponding to matches or mismatches, while horizontal and vertical moves represent indels and incur a fixed gap penalty (set to 0 in this scenario). DTW, on the other hand, adds the residue-residue similarity *S*_*ij*_ at every position visited by the warping path, regardless of whether that position is reached by a diagonal, horizontal, or vertical move. Consequently, DTW requires no explicit gap penalty and produces no explicit gaps. Instead, DTW permits one amino acid to map to multiple contiguous resides in the other sequence. These correspondences are subsequently disambiguated to infer gaps (see **Gap inference**).

### Constructing MSAs from pairwise alignments with template sequence

Building on the pairwise DTW alignment primitive above, we extend our pipeline to the MSA setting. Let 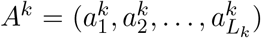 be the *k*^th^ sequence in a protein family with *N* sequences, and 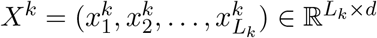 be its PLM embeddings. The conventional strategy for building an MSA is to employ a progressive alignment scheme, where pairwise alignments are merged hierarchically along a guide tree (Thompson et al. 1994; Sievers et al. 2011). However, in our framework,

DTW requires embeddings as input and thus one must construct an embedding-level representation for each intermediate aligned cluster before proceeding to the next merge. This necessitates repeatedly committing to an alignment between a subset of sequences and forming a corresponding consensus embedding. As with progressive alignments within traditional MSA approaches, errors introduced early in the process are amplified in subsequent steps, potentially degrading the overall alignment quality. This is a well-known issue of progressive alignment, and is addressed specifically in methods such as T-Coffee (Notredame et al. 2000) and ProbCons (Do et al. 2005).

To circumvent the issue of repeatedly generating embeddings for intermediate alignments, we instead adopt a two-phase star-alignment strategy, in which a single template is synthesized from the *K* most central sequences, and then this template serves as a common reference frame to align all sequences. By anchoring each DTW alignment to the same coordinate system, we prevent the propagation of incorrect correspondences across multiple pairwise alignments (see Appendix E, Fig. S13, for a comparison between our star-alignment approach and some progressive variants). In what follows, we detail our strategies for merging the resulting set of pairwise DTW alignments into a coherent MSA, and constructing the template sequence to maximize the effectiveness of the star alignment scheme.

#### Star alignment procedure

Our star-alignment scheme can be described independently of the procedure used to construct the template sequence. Let this arbitrary template sequence be of length *L*_*τ*_, and let 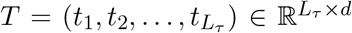 denote the embeddings of its amino acids. Note that *T* may be constructed directly in the embedding space without requiring a semantically valid protein sequence. We further let 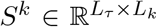 denote the pairwise similarity matrix between the template sequence and the *k*^th^ sequence, and *P* ^*k*^ denote the DTW warping path obtained for this pair. Without loss of generality, we assume that for each (*i, j*) ∈ *P* ^*k*^, *i* refers to the index of the template sequence.

#### Gap inference

To construct a coherent MSA from the set of warping paths 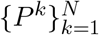, we iterate over each template position *i* and generate a corresponding column block *Q*_*i*_ for the final alignment. Each sequence *X*^*k*^ then contributes to *Q*_*i*_ all residues that associate with *t*_*i*_ in the warping path 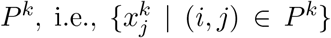 forming the column block’s *k*^th^ row. As DTW permits multi-to-one assignments, a single column block may receive several contiguous residues from the same sequence. Conversely, a single residue in a sequence may be assigned to multiple column blocks. To resolve ambiguities in column assignment, whenever a residue is mapped to multiple column blocks, we assign it the most likely column based on its embedding similarity scores with the template residues.

Letting 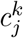 denote the column block that 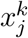 is assigned to:

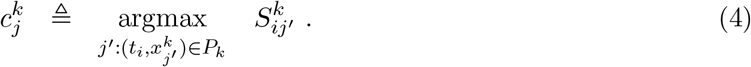

Following this disambiguation step, the *k*^th^ row of column block *Q*_*i*_ is given by 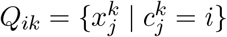. By construction, all column blocks are mutually disjoint across the template positions. However, not all of them are trivially aligned because each sequence can contribute a different number of residues to the same column block. In order to obtain a coherent alignment, we need to infer appropriate gap placements within each block. Our strategy is to identify an anchor residue for each row *Q*_*ik*_ that acts as a true alignment point between *X*_*k*_ and the template. The remaining residues on *Q*_*ik*_ will be positioned on either side of this point, forming a contiguous stretch to avoid fragmented gaps. Finally, we add the required number of gaps to both ends of each stretch to produce a cleanly aligned column block.

In particular, the anchor residue on row *Q*_*ik*_ is again determined by comparing embedding similarity scores. Letting 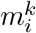 denote the index in *X*^*k*^ that is inferred to align to *t*_*i*_, we have

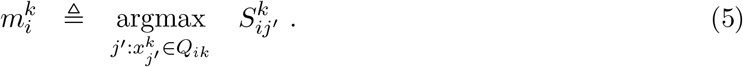

Residues preceding 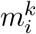 in *Q*_*ik*_ are treated as a single, contiguous insertion to the left of the alignment column, and are appended to the left of the anchor column. Analogously, residues following *m*^*k*^ in *Q*_*ik*_ are treated as a single, contiguous insertion and are appended to the right of the anchor column. Our MSA construction procedure is detailed in Alg. S1, Appendix A.

### Constructing the template sequence

To obtain a representative template sequence for our star-alignment procedure, we first construct a guide tree using Clustal Omega’s mBed algorithm (Sievers et al. 2011). The key idea behind mBed is to avoid computing all *N* ^2^ pairwise distances. Instead, it selects log *N* random anchor sequences from the input set and encodes each sequence as a vector of its distances to these anchors. The distance between any two sequences can then be implicitly approximated by the Euclidean distance between their encoded vectors, enabling construction of a distance-based tree in *O*(*N* log *N* ) time (Sokal et al. 1958).

From this tree, we select the medoid sequence (i.e., the leaf with minimum total distance to all other leaf nodes) as the star-alignment template. We compute this efficiently via an *O*(*N* ) rerooting dynamic programming procedure that performs two depth-first traversals (Sleator and Tarjan 1981): the first pass computes, for each node, the sum of distances to its descendant leaves; the second recursively “reroots” the tree, updating each child’s total distance based on how distances change when the root shifts across an edge (see Alg. S2 in Appendix A).

For larger sequence sets, a single medoid can bias the alignment, especially when sequences are highly divergent or span multiple remote subfamilies; this is a known limitation of standard star-alignment, which assumes one representative sequence can anchor all others. To mitigate this, we construct a PLM-generated consensus template that integrates information from multiple medoids rather than relying on a single one.

We select the top-*K* medoids using the same procedure as before at no extra cost, then align these medoids using our star-alignment method (with the top-1 medoid as the initial template). Gap characters in this MSA are replaced with ‘X’ tokens, and the modified aligned medoids are passed through the PLM to obtain updated embeddings. We then average embedding vectors position-wise across the *K* aligned sequences to form a consensus template, which is used to align the full input set. Empirically, *K* = ⌈ln(*N* )⌉ works well and adds negligible runtime. The pseudocodes of ARIES template selection and overall ARIES multiple sequence alignment are respectively given in Alg. S3 and S4, Appendix A. We further ablate the effect of varying *K* in **Ablation studies**.

### Benchmarking

#### Datasets

We evaluate ARIES on BAliBASE 3.0 (Thompson et al. 2005), HOMSTRAD (Stebbings and Mizuguchi 2004), and QuanTest2 (Sievers and Higgins 2019). BAliBASE is built from structurally-derived reference alignments and defines “core blocks” that identify reliably aligned regions; we evaluate performance on these core blocks. BAliBASE is divided into sequence sets covering diverse challenging scenarios such as low sequence identity, divergent outliers, remote subfamilies, and sequences with large terminal or internal extensions. Accuracy is evaluated with respect to these core blocks. HOMSTRAD consists of protein families whose alignments are based on structural superposition, making it a reliable gold standard for assessing accuracy within conserved regions. QuanTest2 includes sequence sets composed of three HOMSTRAD reference sequences with known structures, augmented with 997 background sequences obtained from Pfam (Mistry et al. 2021); we align all 1,000 sequences but evaluate accuracy only on the reference trio, enabling large-scale assessment while retaining structural ground truth.

Our evaluation consists of 218 BAliBASE alignments, 571 HOMSTRAD families, and 151 Quan-Test2 sets. In some cases, other methods do not output an alignment for some sets; if this happens, we ignore these sets when comparing the method to ARIES. Further information about these three benchmarking datasets are given in Appendix C.

#### Parameter settings

For ARIES, we use a default window span *w* = 9 (or window size 19), reciprocal weight *λ* = 400, and sequence embeddings formed by concatenating outputs from the last *l* = 9 layers of the 650M parameter ESM-2 model (Lin et al. 2023). The star alignment template is generated using the top ⌈ln *N* ⌉ medoids. We performed ablation studies to analyze the effect of these parameters (see **Ablation studies**). For sequences that are longer than 1022, the maximum amino acid context length seen during training of ESM-2, ARIES uses a tiling approach (Brandes et al. 2023) to approximate the amino acid embeddings (see description and performance analysis in Appendix E, Fig. S9). This affects 46 alignment sets in BAliBASE benchmark, and none of the alignment sets in the other two benchmarks.

Clustal Omega (Sievers et al. 2011), ClustalW (Thompson et al. 1994), MAFFT Auto, MAFFT L-INS-i, MAFFT G-INS-i (Katoh and Standley 2013), MAGUS (Smirnov and Warnow 2021), TWI-LIGHT (Tseng et al. 2025), MUSCLE (Edgar 2004), ProbCons (Do et al. 2005), T-Coffee (Notredame et al. 2000), and FAMSA (Deorowicz et al. 2016) are run on CPU with their default parameters. Methods that allow the specification of substitution matrices are evaluated using the default BLOSUM-62 matrix. See Appendix E, Fig. S10, S11, S12 for ablation studies examining the relative performance of alternative substitution matrices. The PLM-based methods learnMSA2 (Becker and Stanke 2024) and vcMSA (McWhite et al. 2023) are run on GPU with recommended default settings. We use ProtT5-XL-Half for learnMSA2 as it is the default PLM for that method. For vcMSA, we pad 10 ‘X’ tokens at each sequence beginning/end and use the last 16 layers of ProtT5-XL. To ensure a fair comparison where PLM-based baselines use the same underlying model, we conduct additional experiments comparing ARIES with ProtT5-XL-Half to learnMSA2, and ARIES with ProtT5-XL to vcMSA (see Appendix E, Table S4). Details of the employed baselines and PLMs are given in Appendices B and C.

#### Evaluation metrics

We evaluate alignments using the Total Column (TC) and Sum-of-Pairs (SP) scores. The TC score is the fraction of columns in the output alignment that exactly match columns in the gold-standard reference alignment, while the SP score is the fraction of residue pairs in the output that are also paired in the reference. Formal definitions of these metrics are provided in Eq. (S2) and Eq. (S1) in Appendix C.

#### Runtime evaluation

For our runtime evaluations on the QuanTest2 dataset, each benchmark is allotted 10 CPU cores with 10 GB of memory. GPU-accelerated methods (ARIES and learnMSA2) are run on a single NVIDIA A100 GPU (80 GB) with the same CPU and memory allocation.

## Results

### Reciprocal, window-based embedding similarity enables detection of aligned residues

We assess our similarity scores (see **ARIES similarity score matrix**) under low-identity conditions with sequence pairs from the BAliBASE RV11 subset (i.e., alignments within BAliBASE with the lowest average pairwise sequence identity). The left panel of Fig. 2 compares the gold-standard alignment of one pair (top-left heatmap) against its pairwise BLOSUM-derived similarity matrix and four embedding-based scoring variants: (a) NED; (b) windowed NED (*w* = 9); (c) reciprocal-weighted NED (*λ* = 400); and (d) ARIES scoring scheme with our default parameters (*w* = 9, *λ* = 400). All embeddings are obtained using the 650M parameter ESM-2 model (Lin et al. 2023); all similarity scores are normalized to [0, 1].

On the exemplar pair, the BLOSUM matrix, lacking contextual information, shows negligible correlation with the true alignment (Fig. 2, *ρ* = 0.0469). NED by itself increases this 5.8-fold (*ρ* = 0.272). Windowed NED (*ρ* = 0.447) and reciprocal-weighted NED (*ρ* = 0.810) substantially improve correlation, while the ARIES scoring scheme, which combines both, achieves the highest correlation (*ρ* = 0.857). The right panel of Fig. 2 shows Pearson correlation distributions across all RV11 pairs. Trends mirror the exemplar: each augmentation significantly improves correlation over the previous (*p* ≤ 1*e*^*−*4^, one-sided Wilcoxon signed-rank test with Holm-Bonferroni correction). Across alignments, the combination of both augmentations (i.e., ARIES’ design) yields a median correlation 4.57 times greater than that of the original NED score. We study the effects of window size and the reciprocal weight *λ* on alignment quality in a later section (see **Ablation studies**).

### ARIES outperforms existing methods across diverse benchmarks

We compare the alignment accuracy of ARIES against a suite of widely used MSA tools on the BAliBASE and HOMSTRAD benchmarks. We present our results in terms of the SP score metric in Fig. 3 and defer the TC score results to Appendix D, Fig. S2. Volcano plots comparing SP/TC scores for ARIES versus all other methods across all datasets can be found in Appendix D, Fig. S3-S8.

**Fig. 3.**
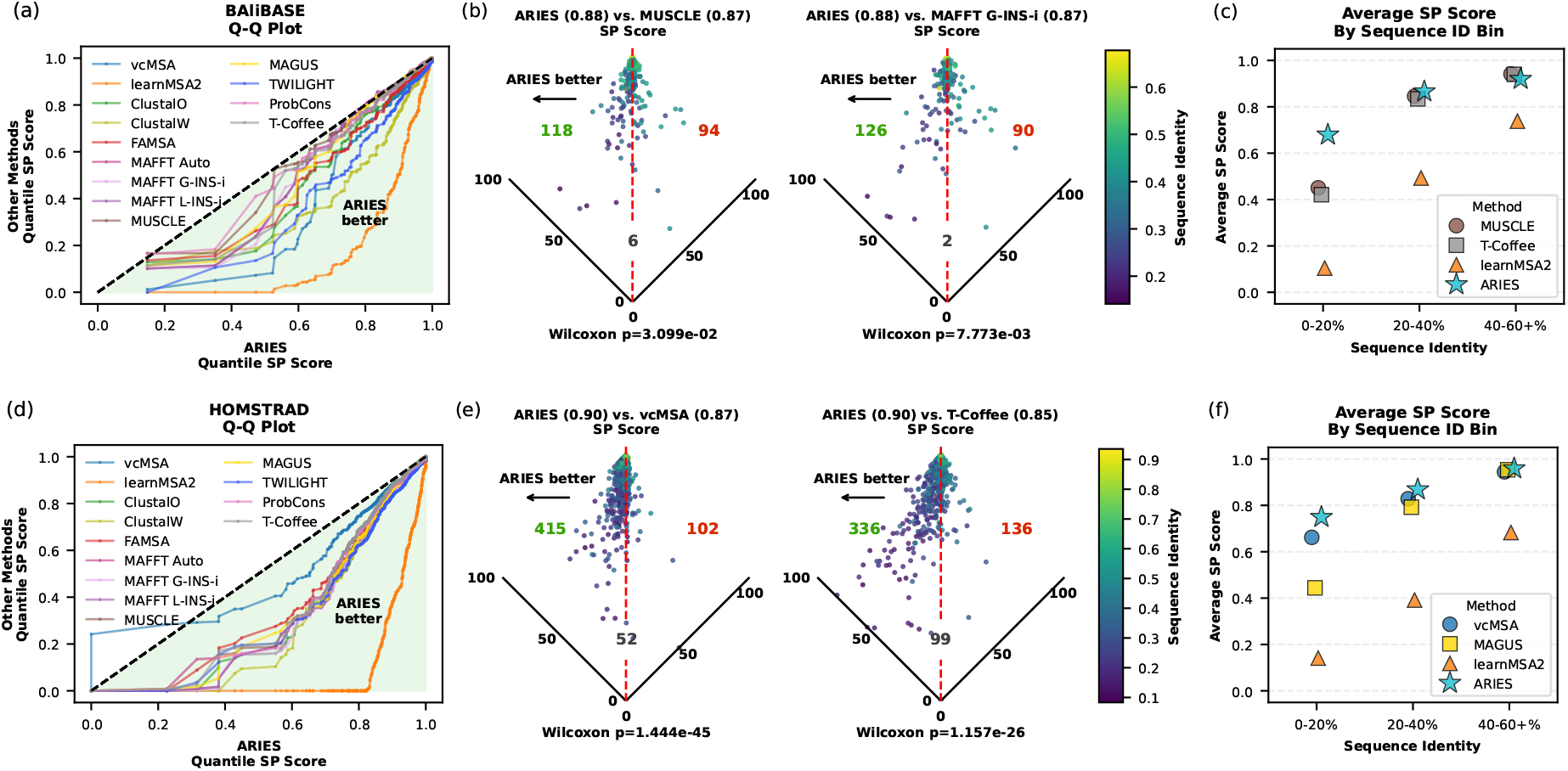
Comparison of ARIES with state-of-the-art MSA methods on (a) BAliBASE and (b) HOMSTRAD. (a, d): Q-Q plots comparing SP scores for ARIES and alternative methods across quantiles; shaded region indicates where ARIES achieves better performance. (b, e): Volcano plots comparing SP scores for ARIES and the two top-performing previous methods, as determined by mean SP score. Each point corresponds to a sequence set that is aligned; those to the left of the vertical line indicate better performance by ARIES. Colored counts show the number of sequence sets for which ARIES performs better than (green), worse than (red), or equal to (gray) the competitor method. Number in bracket next to each method name shows its mean SP score on the entire dataset. Points are colored by mean sequence identity of the MSA set; darker color indicates lower identity. Holm-Bonferroni corrected *p*-values from one-sided Wilcoxon signed-rank tests are annotated at the bottom of each plot. (c, f): Mean SP scores of representative methods in three sequence identity bins.

We showcase the superior performance of ARIES as compared to other methods using Q-Q plots (Fig. 3a,d). Each curve compares ARIES (*x*-axis) to another single method (*y*-axis), with points showing SP scores at the same quantile. The shaded region below the diagonal highlights quantiles where ARIES outperforms the competitor method. Across both datasets, most curves lie below the diagonal, indicating that ARIES consistently exceeds competing methods. On BAliBASE, some methods perform similarly to ARIES at upper quantiles, while lower quantiles—representing more challenging alignments—show pronounced margins, demonstrating ARIES’ robustness where other approaches degrade (see Appendix E, Tables S2, S3 for average SP and TC scores in each of the six BAliBASE subsets). On HOMSTRAD, ARIES exhibits substantially higher SP scores than all other methods, as reflected by curves lying well below the diagonal line of equal performance; vcMSA is second-best, but considerably slower.

Volcano plots (Fig. 3b,e) provide a detailed view of how ARIES compares to the top two performing other methods (as judged by their average SP score across sequence sets in BAliBASE and HOMSTRAD); analogous comparisons for all competing methods are given in Appendix D, Fig. S3-S6. Each point represents a protein family, and is plotted based on its SP score for ARIES (left axis) and the competing method (right axis). Points to the left of the vertical line represent cases where ARIES produces more accurate alignments. Across all comparisons (Appendix D, Fig. S3), the plots show a strong leftward skew, with far more points favoring ARIES than the other methods. On BAliBASE, ARIES outperforms the next-best approach, MUSCLE, on 54.1% of sequence sets. On HOMSTRAD, ARIES performs better than vcMSA, the next best method, in 72.9% of cases. Against the worst performing baseline, learnMSA2, ARIES outperforms it in more than 85% sequence sets on both datasets (see Appendix D, Fig. S3, S5). Using one-sided Wilcoxon signed-rank tests with Holm–Bonferroni correction, we confirm that ARIES’ SP score improvements over all competing baselines are statistically significant on BAliBASE (largest corrected *p* = 0.031), and HOMSTRAD (largest corrected *p* = 1.157*e*^*−*26^). The TC score improvements over other baselines are all statistically significant (see volcano plots for TC scores in Appendix D, Fig. S4, S6, largest corrected values of *p* = 0.024 and *p* = 1.454*e*^*−*22^ on BAliBASE and HOMSTRAD, respectively).

Further, across both datasets, the alignment sets with lowest sequence identity (depicted with darker colors) tend to fall to the left of the vertical line, reinforcing that ARIES performs particularly well in this “twilight zone” of very low sequence identity. We quantify this trend by evaluating the average SP scores of ARIES and three representative baselines (best, median, and worst performers) on three coarse sequence-identity bins (Fig. 3c,f). ARIES achieves the highest mean SP scores in lower-identity bins (0–20% and 20–40% on HOMSTRAD, and 0-20% on BAliBASE) and matches the top and median methods otherwise. In contrast, learnMSA2, which was designed to align large sequence families, consistently performs worst across all bins, suggesting it is indeed poorly suited for the small-to medium-sized families in BAliBASE and HOMSTRAD. We also break down the performance of all methods across the six BAliBASE subsets. ARIES’ performance relative to the baselines is strongest on RV11, which has the lowest overall sequence identity (see Table S2 and S3).

### ARIES scales efficiently and accurately to large datasets

We further evaluate ARIES in large-scale settings using the QuanTest2 benchmark, which contains 151 alignments with 1000 sequences each. We compare ARIES to the nine baseline approaches that scale well to this input size, defined as those taking less than 10 minutes per set on average with default parameters (Fig. 4a). Volcano plots for all pairwise baseline comparisons are given in Appendix D, Fig. S7 (SP score) and Fig. S8 (TC score).

**Fig. 4.**
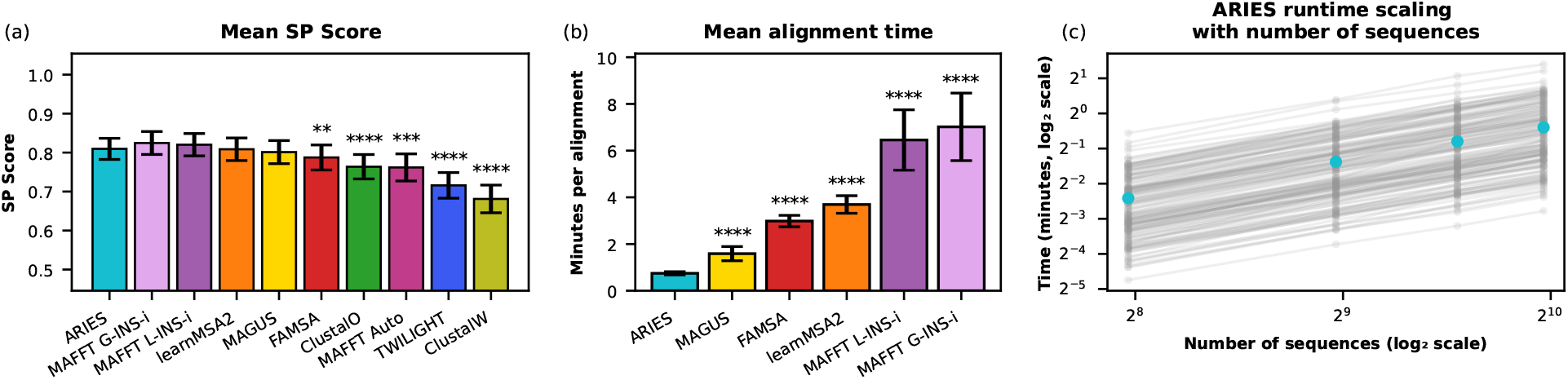
Comparison of ARIES with other MSA methods on the QuanTest2 dataset. (a) For each scalable method considered, the mean SP score across all QuanTest2 alignments, shown with 95% confidence intervals. (b) Mean alignment runtime (minutes per alignment) for methods that ARIES does not significantly outperform, shown with 95% confidence intervals. ARIES has the lowest mean runtime of these methods, and its runtime is significantly lower than that of all of these methods. Stars in panels (a) and (b) denote Holm-Bonferroni corrected one-sided paired Wilcoxon signed-rank tests across alignments (^*\**^ *p ≤* 0.05, ^*\*\**^ *p ≤* 1*e*^*−*2, *\*\*\**^ *p ≤* 1*e*^*−*3, *\*\*\*\**^ *p ≤* 1*e*^*−*4^). (c) log-log plot of ARIES runtimes for each QuanTest2 family across varying subset sizes. Lines show individual families, with blue points indicating the mean runtime at each size tested.

ARIES obtains higher mean SP scores than all methods other than MAFFT G-INSI and MAFFT L-INSI. However, the two MAFFT variants do not perform significantly better than ARIES (Fig. S7, one-sided Wilcoxon Holm-Bonferroni corrected *p*-values of 0.0949 and 0.142, respectively), and likewise ARIES does not significantly outperform learnMSA2 and MAGUS (the latter of which merges alignments obtained by MAFFT L-INSI). On the other hand, ARIES significantly outperfoms ClustalO, ClustalW, FAMSA, TWILIGHT and MAFFT Auto (Fig. S7, one-sided Wilcoxon test with Holm-Bonferroni correction).

ARIES provides substantial runtime improvements as compared to the other MSA methods whose performances are neither significantly worse or significantly better than it (Fig. 4b). Within this set, ARIES is significantly faster than MAGUS, FAMSA, learnMSA2, MAFFT L-INS-i, and MAFFT G-INS-i under the same resource budget (see **Benchmarking** for hardware description). We note that both ARIES and learnMSA2 are GPU-accelerated, and thus most directly comparable, while the other methods run on CPU only. Full runtime results are in Fig. S15.

To further evaluate scaling, we measure runtime of ARIES on progressively larger subsets of each family in QuanTest2 (250, 500, 750, and 1000 sequences). Fig. 4c shows a log-log plot of the runtime (in minutes) versus sequence count, revealing approximately linear scaling with the number of sequences. The slope in log-scale is 1.010 (95% confidence interval: 0.920, 1.110), consistent with approximately linear scaling. In theory, our approach can scale even further with multi-GPU parallelism and larger accelerator clusters. This establishes ARIES as a highly efficient PLM-based alignment approach for large sequence families.

### Ablation studies

We next examine how ARIES’ hyperparameters affect performance. Specifically, we conduct controlled experiments to probe the impact of (a) window and reciprocal-weight parameters in the similarity metric, (b) embedding depth across different PLM backbones, and (c) the number of medoids used for template construction. Results of these ablations are shown in Fig. 5a–c, with experimental details provided below.

**Fig. 5.**
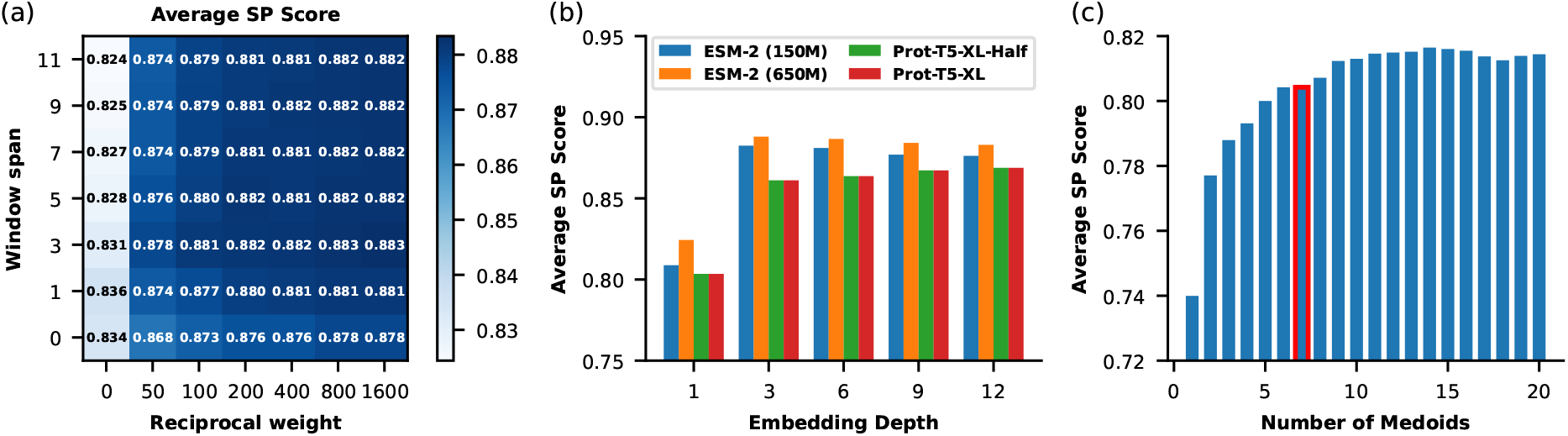
Ablation study of key components of ARIES. (a) Average SP score on BAliBASE dataset across 49 combinations of window sizes and reciprocal-weight parameters; (b) Average SP score on BAliBASE dataset across varying embedding depth using four different PLM backbones; and (c) Average SP score on QuanTest 2.0 dataset with varying number *K* of medoid sequences used in the template-building stage. The red-outlined bar marks *K* = *⌈*ln(*N* )*⌉*.

#### Effect of varying window and reciprocal-weighting parameters

Fig. 5a presents a heatmap summarizing SP scores on the BAliBASE dataset across 49 hyperparameter settings. We vary the window span *w* ∈ {0, 1, 3, 5, 7, 9, 11}, corresponding to window lengths in {1, 3, 7, 11, 15, 19, 23}), and the reciprocal weight parameter *λ* ∈ {0, 50, 100, 200, 400, 800, 1600}. Here, setting *w* = 0 and *λ* = 0 corresponds to using just NED. Across these combinations, we observe a significant jump in SP score moving from a residue-based metric (*w* = 0) to any window-based metric (*w* ≥ 1) at all *λ* values (one-sided Wilcoxon *p*-values ≤ 1.27 × 10^*−*5^ for each of the seven comparisons). This pattern demonstrates that incorporating local context indeed improves the robustness of alignment signals, which agrees with our earlier finding on metric fitness (see **Reciprocal, window-based embedding similarity enables detection of aligned residues**). Performance remains relatively consistent as we further increase *w*. We attribute this stability to the smoothing effect of the discrete Gaussian weights: residues further away from the center naturally receive lower influence, allowing the method to maintain robustness even as the window size increases. We also find that increasing *λ* consistently leads to meaningful improvements in alignment quality, with improvements saturating by *λ* = 800, confirming that mutual agreement between residues provides a robust and reliable alignment signal.

#### Choice of PLM and embedding depth

Fig. 5b examines the effect of embedding depth on alignment quality using four PLM backbones: ESM-2 (150M), ESM-2 (650M) (Lin et al. 2023), ProtT5-XL-Half, and ProtT5-XL (Elnaggar et al. 2021). For each model, ARIES SP scores on BAliBASE are computed when representing amino acids using concatenations of the last *l* PLM layer embeddings, with *l* ∈ [1, 3, 6, 9, 12]. Across all models, the largest gain occurs from *l* = 1 to *l* = 3, likely because the final layer emphasizes residue identity over structural context. Runtime increases with embedding dimension (see Appendix D, Fig. S16). Relative PLM rankings remain consistent, with ESM-2 (650M) performing best, indicating ARIES is model-agnostic yet benefits from stronger embeddings.

#### Template sequence construction

Fig. 5c illustrates how the number of medoid sequences *K* influences the template quality on the QuanTest2 dataset. Incorporating more medoid sequences naturally allows the template to incorporate patterns from a broader range of structural subgroups, which is especially beneficial for large and diverse families such as those in QuanTest2. As expected, ARIES shows sharp improvement in SP scores going from one medoid sequence to larger values of *K*, which demonstrates the weakness of a simple star-alignment stragety and the importance of well-informed template generation. The improvements seem to saturate beyond *K* = 14, which suggests that a relatively small number of representative sequences already capture most of the diversity in the MSA set. The red-outlined bar highlights the configuration *K* = ⌈ln(*N* )⌉, which achieves near-optimal performance. This empirical setting of *K* also works generally for different MSA set sizes (see Appendix D, Fig. S14). This principled, dataset-adaptive way to set the number of medoids also guarantees an efficient runtime, as it only adds a negligible *O*(log(*N* )) overhead to our pipeline (see empirical runtime analysis in Appendix D, Fig. S15).

## Discussion

We have developed ARIES, a new framework for constructing MSAs that leverages contextual representations from PLMs. ARIES integrates a windowed reciprocal-weighted embedding similarity metric with a DTW-based star-alignment strategy driven by a PLM-generated template, moving beyond the constraints of fixed substitution matrices and explicit gap penalties. Across large-scale evaluations on BAliBASE, HOMSTRAD, and QuantTest2, ARIES consistently delivers substantial, statistically-significant performance gains over state-of-the-art methods while maintaining favorable scaling with growing sequence sets. These results demonstrate that PLM-derived signals can materially advance MSA accuracy for evolutionarily distant proteins and provide a scalable foundation for large-scale alignment.

An important consideration in interpreting these results is the distinction between positional homology, in which residues in the same alignment column are inferred to descend from a common ancestral residue, and structural correspondence, in which residues are aligned based on shared structural position or role regardless of common ancestry. ARIES is agnostic to this distinction: its alignments are driven by PLM embedding similarities, which are learned from sequence but implicitly reflect physicochemical and structural constraints. Because our benchmark datasets are derived from structural superpositions, our reported accuracies primarily measure agreement with structural correspondence. Although this may provide a reasonable proxy for positional homology among closely related sequences, the two can diverge in the twilight zone. Whether PLM-based alignment can reliably recover positional homology for phylogenetic inference therefore remains an important question for future work.

Although ARIES performs well, several directions could further advance embedding-based alignment. Classical MSA ideas, such as iterative refinement (Edgar 2004; Katoh and Standley 2013), consistency transformations (Do et al. 2005; Notredame et al. 2000), and profile–profile alignment (Krogh et al. 1994), could be incorporated to improve accuracy. BLAST-style seeding (Altschul et al. 1990) and related seed selection techniques (Marçais et al. 2017; Hoang et al. 2022, 2024) could also be incorporated to improve runtime scalability. Another opportunity is to learn the similarity metric directly, rather than relying on the zero-shot use of pretrained PLM embeddings; given the abundance of protein families with high-quality MSAs, such training should be feasible. Additionally, recent methods such as SMURF (Petti et al. 2023) and DEDAL (Llinares-López et al. 2023) (Appendix B), although not focused on full MSA construction, highlight the benefits of integrating alignment learning into end-to-end workflows; ARIES could serve as an alignment module in such systems, offering improved performance in low-identity settings and scalability to large families.

Overall, given the central role of MSAs in the structural, evolutionary, and functional analyses of proteins, and the clear improvements in scalability and accuracy demonstrated by ARIES, we anticipate that PLM-based approaches will become a new foundation for large-scale protein alignment.

## Code Availability

ARIES implementation and setup instructions can be found at https://github.com/Singh-Lab/ARIES.

## Acknowledgement

Research reported in this publication was supported in part by National Institute of General Medical Sciences of the National Institutes of Health under grant number R35GM158278, and by the Princeton Laboratory for Artificial Intelligence.

## Author Contribution

M.H., I.A.-G., and M.S.: Conceptualization, Methodology, Writing – original draft. M.H. and I.A.-G.: Software.

## Disclaimer

The content is solely the responsibility of the authors and does not necessarily represent the official views of the National Institutes of Health.

## A ARIES multiple sequence alignment module

Alg. S1 describes the star-alignment procedure (see **Star alignment procedure**). Within this procedure, the subroutine ReciprocalScore implements our embedding-based similarity metric, parameterized by the window size *w* and reciprocal weight *λ* (see **ARIES similarity score matrix**), while DynamicTimeWarping performs the DTW-based pairwise alignment (see **Dynamic Programming for Pairwise Sequence Alignment**).

Alg. S2 details the linear-time rerooting dynamic programming algorithm we use to extract the top-*K* medoids from the Clustal guide tree. This tree is constructed using the mBed method, which embeds sequences into a reduced-dimensional space and clusters them efficiently. The rerooting DP procedure enables us to compute medoid scores at every internal node in *O*(*N* ) time with two depth-first traversals.

Alg. S3 outlines our template generation strategy based on these *K* medoids. After aligning the medoids with our method, we convert their alignment into a generative template by replacing intermediate gaps with ‘X’ tokens, re-embedding the resulting pseudo-sequences, and taking a position-wise average of their embeddings.

Finally, Alg. S4 formalizes the complete ARIES MSA pipeline, integrating medoid extraction, template construction, and star alignment into a unified end-to-end framework.

### Algorithm S1

ARIES Star-Alignment Procedure (StarAlign)

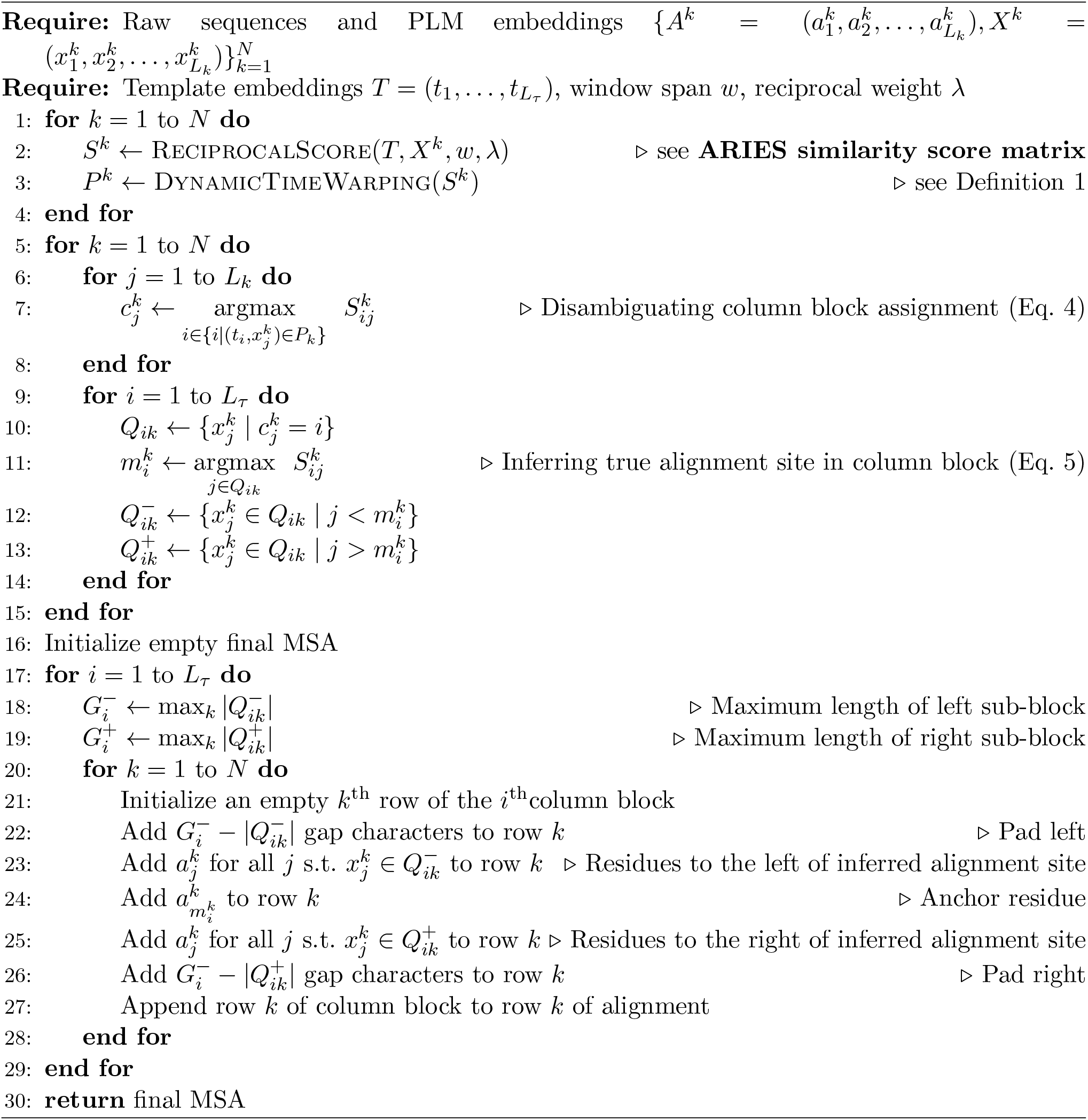

### Algorithm S2

Top-*K* Medoid Selection (Top-K-Medoids)

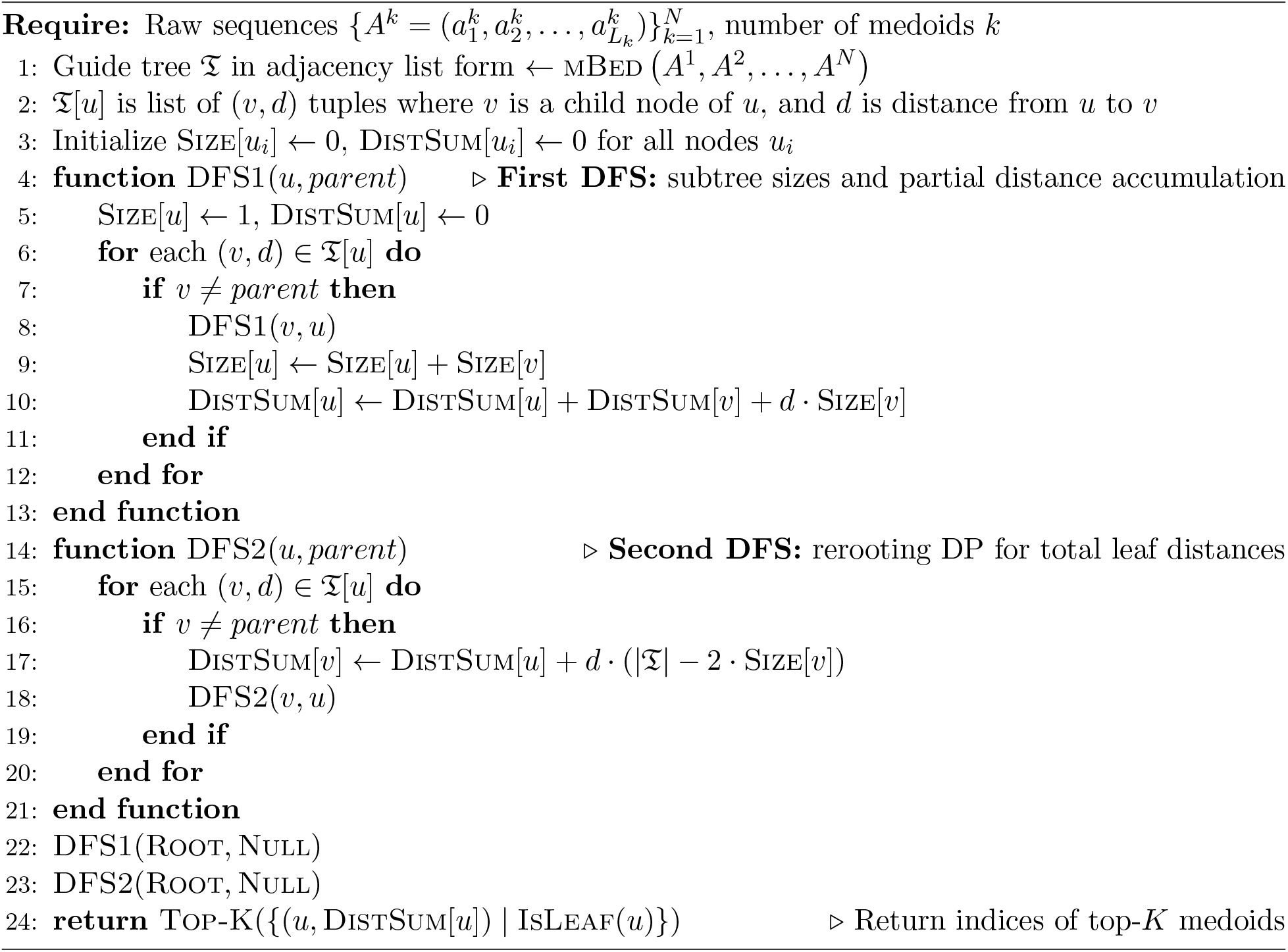

### Algorithm S3

ARIES Template Selection (ConstructTemplate)

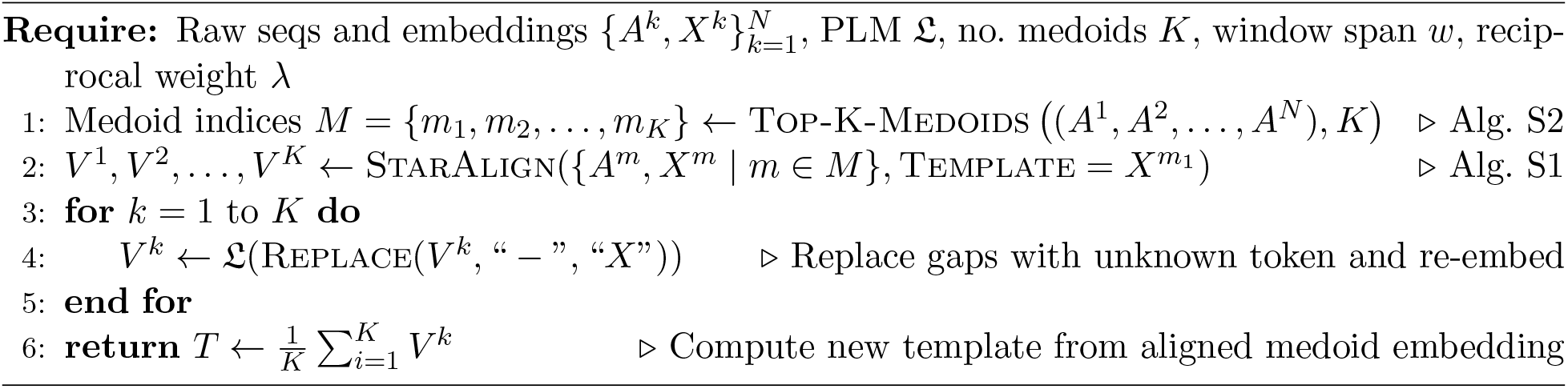

### Algorithm S4

ARIES Multiple Sequence Alignment (ARIES)

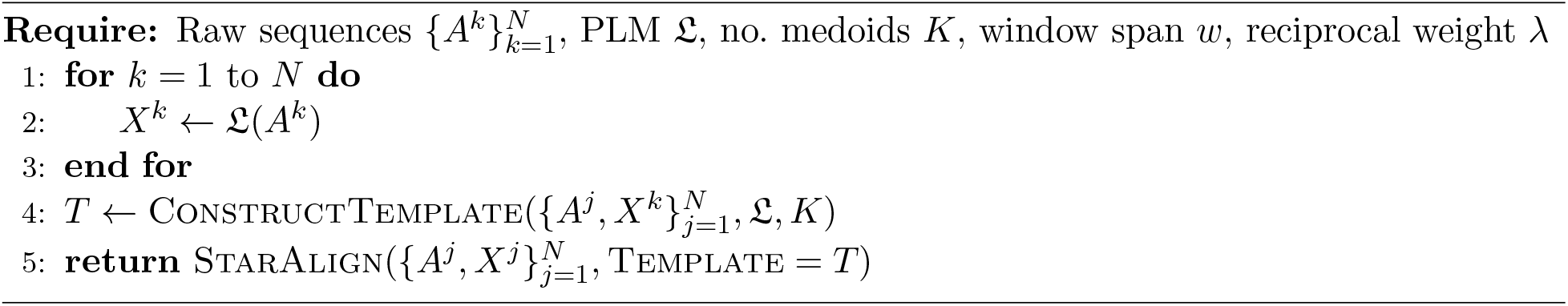

## B Related work

**ClustalW** (Thompson et al. 1994) (version 1.2.4) is a classic progressive alignment algorithm that constructs a guide tree from pairwise distances and performs alignment using position-specific gap penalties and sequence-weighting schemes.

**ClustalO (Clustal Omega)** (Sievers et al. 2011) (version 2.1) is the successor to **ClustalW** and scales to large protein families using mBed sequence embeddings, guide-tree clustering, and optimized progressive alignment.

**MUSCLE** (Edgar 2004) (version 5.1) is an iterative refinement method combining progressive alignment with tree-based realignment cycles, offering a strong balance between computational efficiency and accuracy.

**MAFFT** (Katoh et al. 2002) (version 7.526) is a scalable MSA framework that accelerates pair-wise alignment using fast Fourier transforms and offers multiple iterative refinement strategies for improved accuracy. We evaluate three configurations: **MAFFT L-INS-i**, which performs iterative refinement using local pairwise alignment information, **MAFFT G-INS-i**, which performs iterative refinement using global pairwise alignment information, and **MAFFT Auto**, (the -auto flag), which “automatically selects an appropriate strategy from L-INS-i, FFT-NS-i and FFT-NS-2, according to data size.” For L-INS-i and G-INS-i, we allow up to 1000 refinement iterations.

**T-Coffee** (Notredame et al. 2000) (version 13.46.2) is a consistency-based method that integrates heterogeneous pairwise alignment libraries to guide progressive alignment through a unified scoring matrix.

**FAMSA** (Deorowicz et al. 2016) (version 2.4.1) is a fast progressive alignment tool optimized for very large datasets, using efficient clustering, SIMD-accelerated scoring, and memory-efficient data structures.

**ProbCons** (Do et al. 2005) (version 1.12) is a consistency-based method employing pair-HMM posterior probabilities and maximum expected accuracy optimization.

**TWILIGHT** (Tseng et al. 2025) (version 0.2.3) is a progressive alignment method designed for very large datasets, using parallelization and memory-efficiency strategies.

**MAGUS** (Smirnov and Warnow 2021) (version 0.2.0) is a multiple sequence alignment method for large datasets that aligns sequence subsets independently and merges them through graph-based clustering, rather than relying on traditional guide tree-based progressive alignment.

**vcMSA** (McWhite et al. 2023) (version 1.2.0) is the first PLM-based MSA framework. This method infers MSA through recursive vector clustering. However, the recursive clustering is not scalable for large protein families.

**learnMSA2** (Becker and Stanke 2024) (version 2.0.14) integrates PLM embeddings with a deep HMM architecture. This method is optimized for large protein families, though performance can be unstable on smaller or highly divergent input sets.

**SMURF** (Petti et al. 2023) introduces a differentiable formulation of the Smith–Waterman algorithm that produces a soft alignment tensor approximating a discrete MSA, enabling end-to-end integration with downstream models such as AlphaFold2. **DEDAL** (Llinares-López et al. 2023), **EBA** (Pantolini et al. 2024), and **PEbA** (Iovino and Ye 2024) all leverage **PLM** embeddings to perform pairwise sequence alignment. These methods cannot reconstruct a coherent multiple sequence alignment.

## C Other experiment details

### Protein Language Models

Table S1 below gives the specifications of 4 open-source language models used in our experiments. All PLMs are publicly released on HuggingFace.

**Table S1.** No. parameters, output dimension, and floating point precision of various PLMs used in our experiments.

| PLM | ESM-2 (150M) | ESM-2 (650M) | Prot-T5-XL | Prot-T5-XL-Half |
|---|---|---|---|---|
| No. params | 150M | 650M | 1.2B | 1.2B |
| Output dims | 480 | 640 | 1024 | 1024 |
| Precision | 32 bits | 32 bits | 32 bits | 16 bits |

### Datasets

BAliBASE is a structure-based benchmark comprised of reference alignment sets derived from three-dimensional structural superpositions, followed by manual curation and refinement of the resulting alignments. Each reference set is designed to isolate a specific class of difficult alignment scenarios, including low-identity sequences, highly divergent outliers, remote subfamilies, long terminal expansions, and large internal insertions. These categories (RV11, RV12, RV20, RV30, RV40, and RV50) provide controlled tests of alignment robustness under diverse evolutionary and structural conditions. BAliBASE annotates core blocks, which are subsets of alignment columns corresponding to structurally conserved regions that are reliably aligned. Performance is assessed with respect to these core blocks.

HOMSTRAD (Homologous Protein Structure Alignment Database) is comprised of protein families grouped by sequence and structural similarity, for which high-quality experimentally-determined structures are available. For each family, alignments are generated using structural superposition rather than sequence similarity alone, and are then curated to ensure consistency within conserved structural regions. This construction makes HOMSTRAD a reliable benchmark for evaluating whether sequence-based methods recover structurally-meaningful residue correspondences in well-characterized protein folds.

QuanTest2 is a large-scale structure-based benchmark designed to assess alignment performance in high-depth and potentially noisy settings. Each alignment contains exactly 1000 sequences: three reference sequences drawn from HOMSTRAD with known three-dimensional structures, supplemented with 997 additional homologous sequences sampled from Pfam to span broader sequence diversity. Methods are applied to all 1000 sequences, but accuracy is evaluated only on the induced alignment of the three reference sequences. This design enables large-scale testing on diverse sequence collections while retaining a reliable structural ground truth for scoring.

Fig. S1 shows several key statistics of the three datasets used throughout our empirical studies. Across three datasets, HOMSTRAD has the smallest input size (on average 3.52±3.72 sequences per set), followed by BAliBASE (28.54 ± 26.23 sequences per set), whereas every set in the QuanTest2 dataset has exactly 1000 sequences. BAliBASE has both longest maximum sequence length and largest sequence length variance among all 3 datasets, while QuanTest2 has the lowest average sequence identity.

**Fig. S1.**
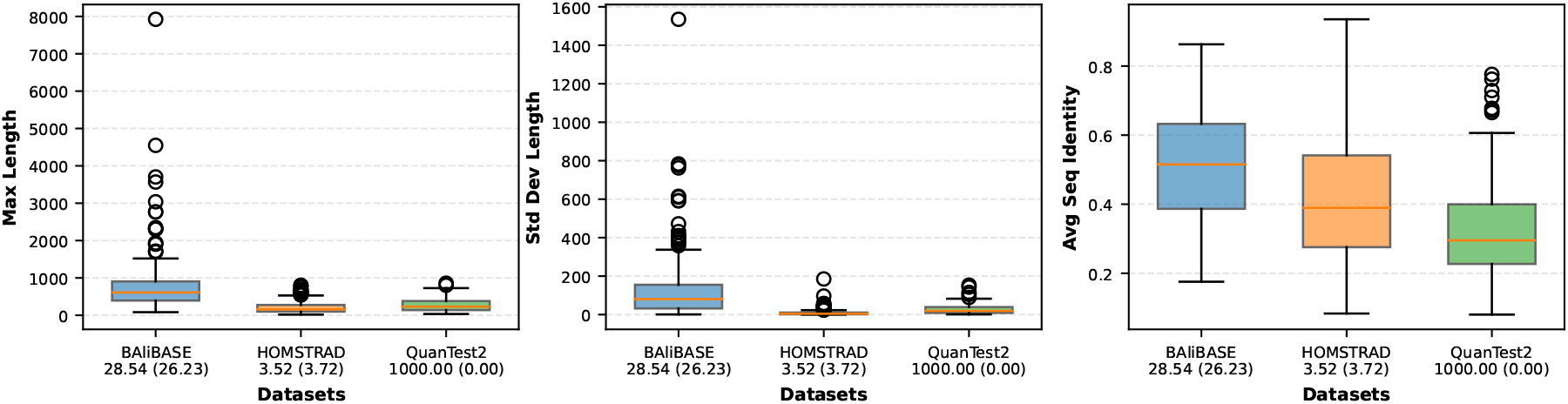
Key statistics of the BAliBASE, HOMSTRAD, and QuanTest2 datasets. We report the mean number of sequences per family, along with its standard deviation (in parentheses), beneath each dataset label on the x-axis. *Left:* Box plots of maximum sequence length in each protein family. *Middle:* Box plots of standard deviation of sequence lengths in each protein family. *Right:* Box plots of average sequence identity in each protein family. For QuanTest2 we measured the average sequence identity among the three reference sequences.

### Metrics

Let ℛ be the set of tuples (*i, j, u, v*) specifying that residue *u* of sequence *i* aligns with residue *v* in sequence *j* in the reference alignment, and *P* be the corresponding set for the predicted alignment. The **Sum-of-Pairs (SP) score** is defined as:

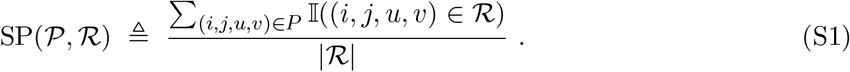

Further let *L*_*r*_ and *L*_*p*_ be the number of columns in ℛ and *P*; *P*_*c*_ be the set of all tuples (*i, u*) specifying that residue *u* of sequence *i* is assigned to column *c* of *P* ; and col(*R, i, u*) denote the column in *R* that residue *u* of sequence *i* is assigned to. The **Total Column (TC) score** is defined as

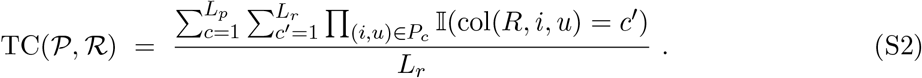

## D Additional evaluation of ARIES

### ARIES achieves better SP and TC scores than existing methods across diverse benchmarks

To complement the summarized SP results reported in the main text, we additionally provide the summarized TC results (Fig. S2) and the complete set of volcano plots comparing ARIES against all other baselines with respect to both SP and TC scores (Figs. S3 and S8).

**Fig. S2.**
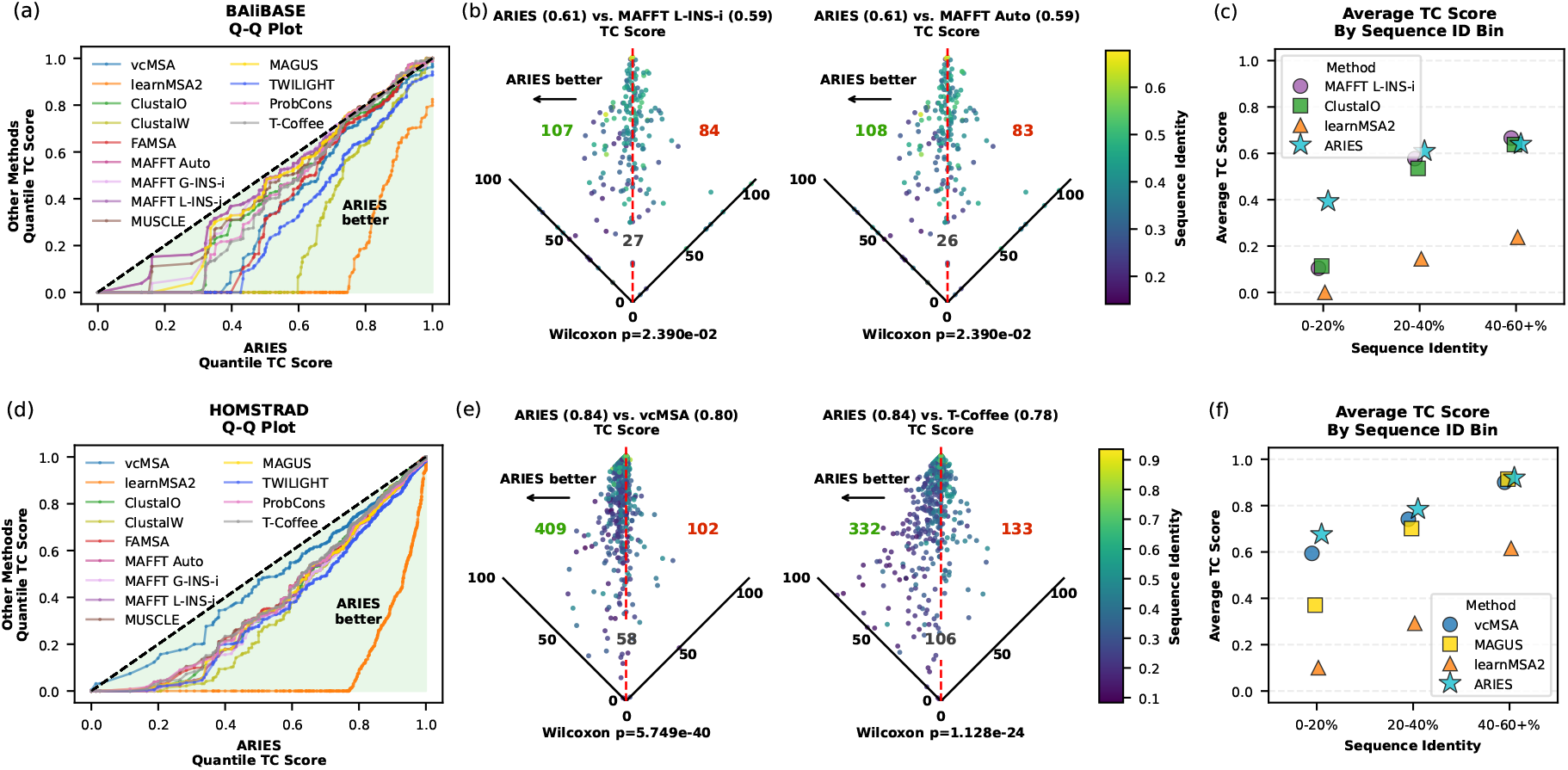
Comparison of ARIES with state-of-the-art MSA methods on (a) BAliBASE and (b) HOMSTRAD. (a, d): Q-Q plots comparing TC scores for ARIES and alternative methods across quantiles; shaded region indicates where ARIES achieves better performance. (b, e): Volcano plots comparing TC scores for ARIES and the two top-performing previous methods, as determined by mean TC score. Each point corresponds to a sequence set that is aligned; those to the left of the vertical line indicate better performance by ARIES. Colored counts show the number of sequence sets for which ARIES performs better than (green), worse than (red), or equal to (gray) the competitor method. Number in bracket next to each method name shows its mean TC score on the entire dataset. Points are colored by mean sequence identity of the MSA set; darker color indicates lower identity. Holm-Bonferroni corrected *p*-values from one-sided Wilcoxon signed-rank tests are annotated at the bottom of each plot. (c, f): Mean TC scores of representative methods in three sequence identity bins.

### Head-to-head SP score comparison with all baselines on BAliBASE dataset

**Fig. S3.**
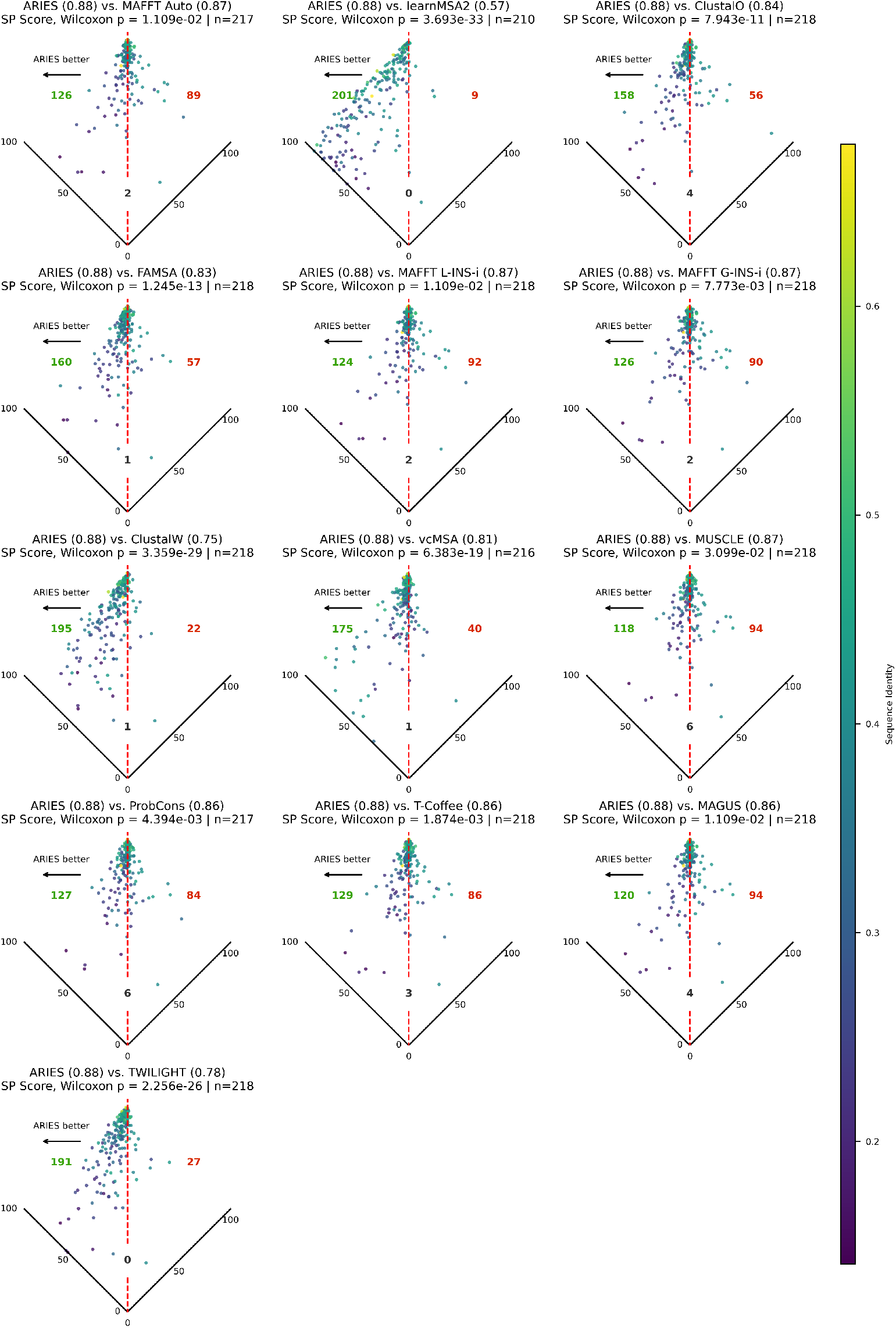
Comparison of ARIES with state-of-the-art MSA methods on BAliBASE using SP scores. Mean SP scores over the entire dataset are given next to method names; Holm-Bonferroni-corrected *p*-value of one-sided Wilcoxon signed-rank test is reported for each baseline method.

### Head-to-head TC score comparison with all baselines on BAliBASE dataset

**Fig. S4.**
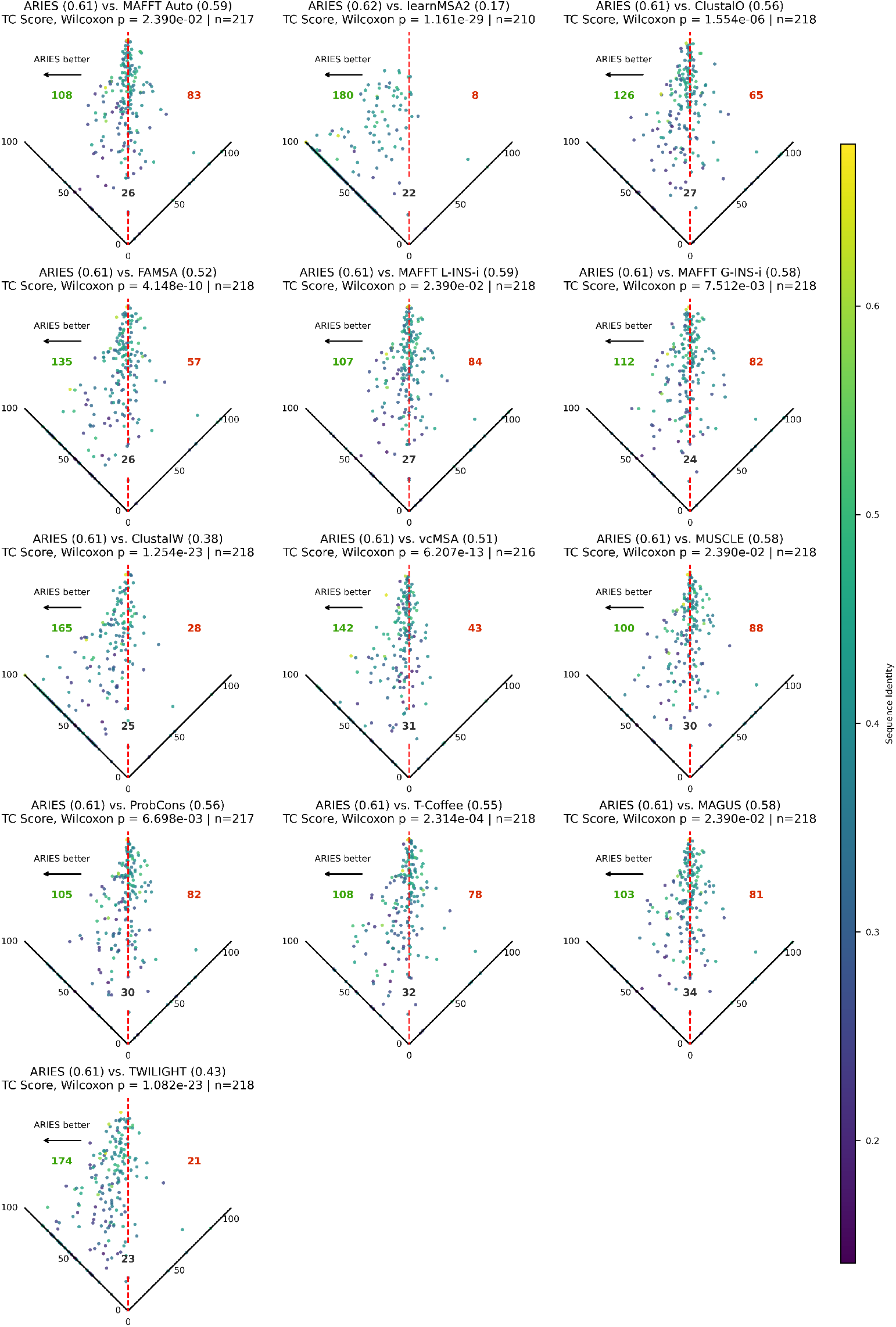
Comparison of ARIES with state-of-the-art MSA methods on BAliBASE using TC scores. Mean TC scores over the entire dataset are given next to method names; Holm-Bonferroni-corrected *p*-value of one-sided Wilcoxon signed-rank test is reported for each baseline method.

### Head-to-head SP score comparison with all baselines on HOMSTRAD dataset

**Fig. S5.**
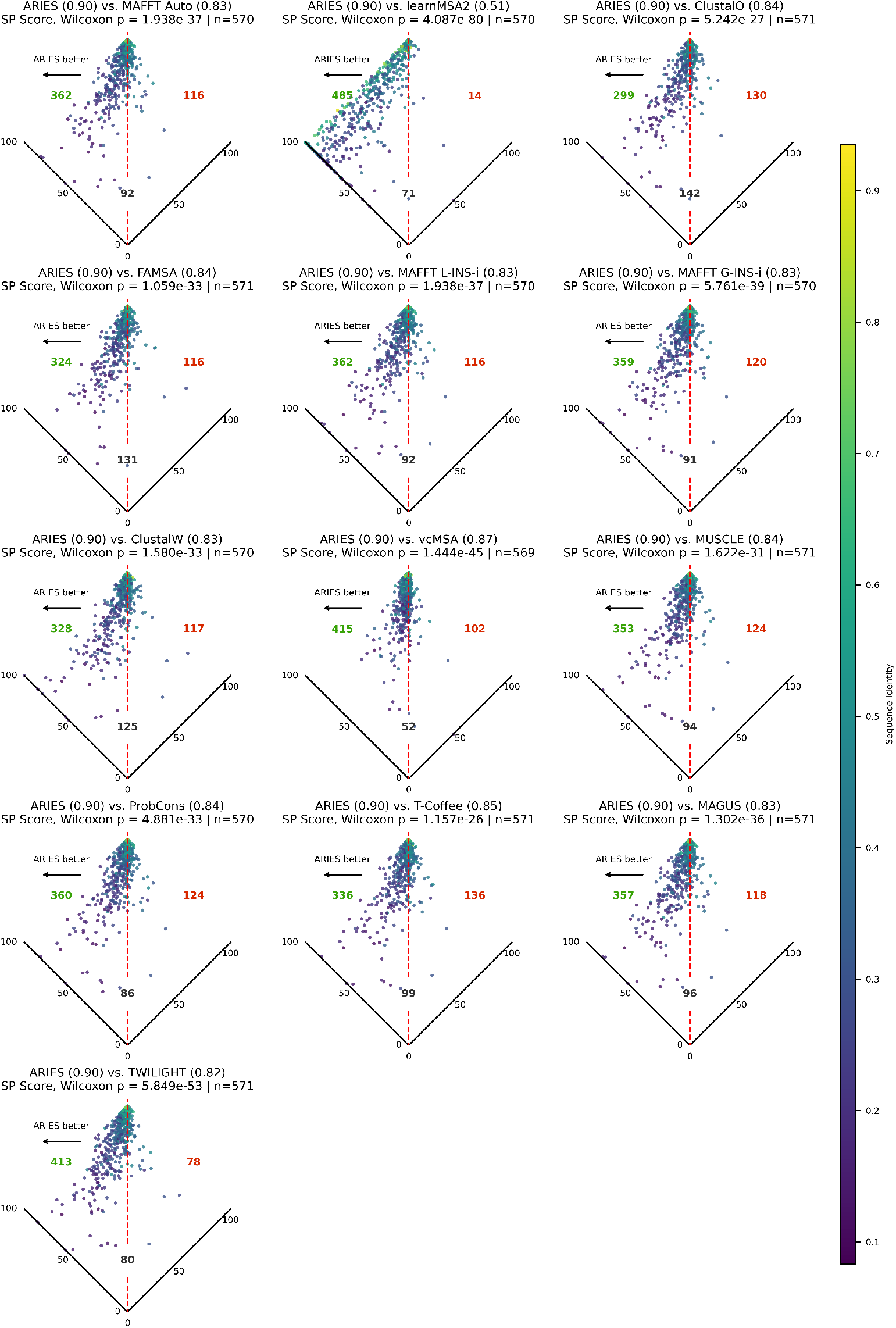
Comparison of ARIES with state-of-the-art MSA methods on HOMSTRAD using SP scores. Mean SP scores over the entire dataset are given next to method names; Holm-Bonferroni-corrected *p*-value of one-sided Wilcoxon signed-rank test is reported for each baseline method.

### Head-to-head TC score comparison with all baselines on HOMSTRAD dataset

**Fig. S6.**
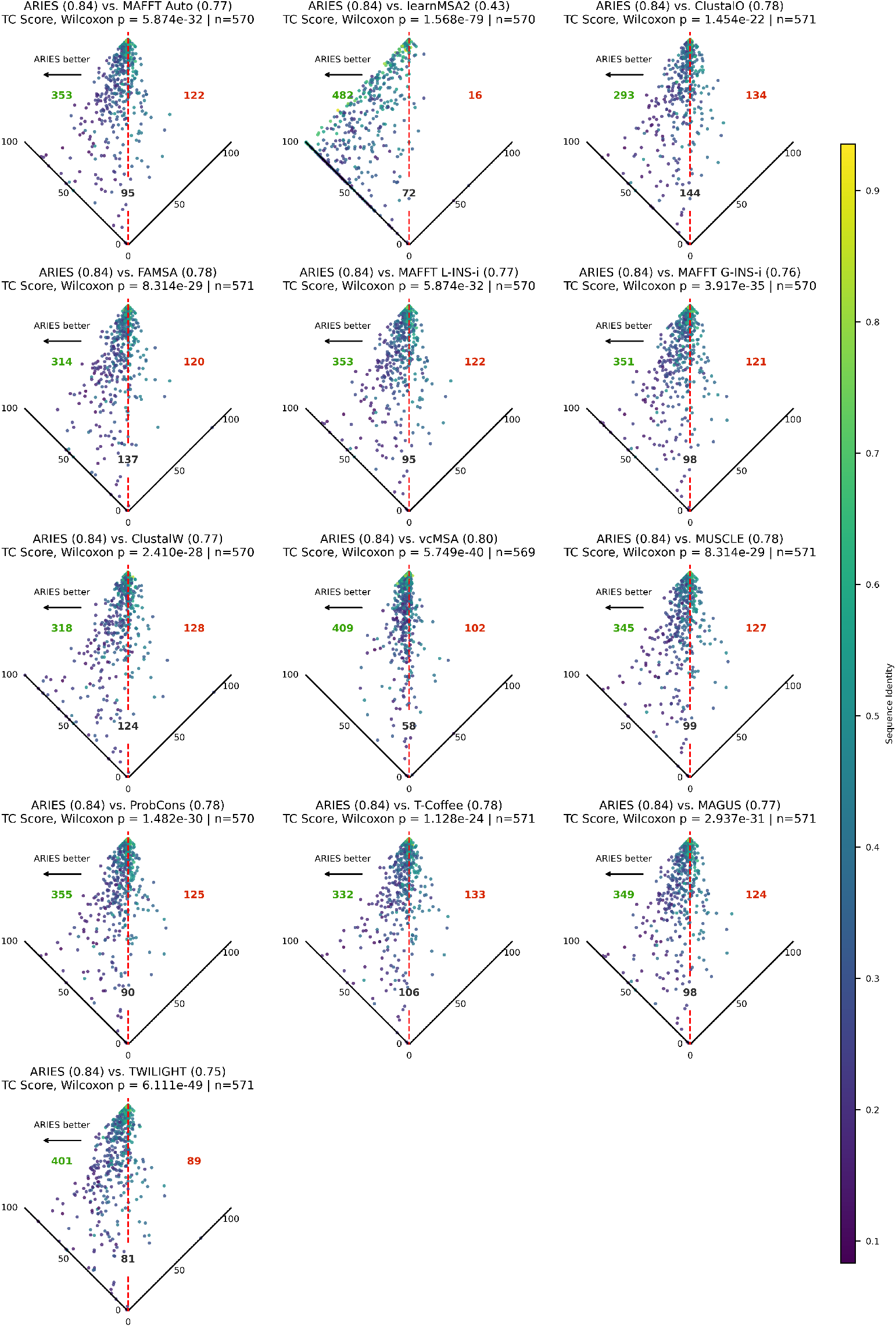
Comparison of ARIES with state-of-the-art MSA methods on HOMSTRAD using TC scores. Mean TC scores over the entire dataset are given next to method names; Holm-Bonferroni-corrected *p*-value of one-sided Wilcoxon signed-rank test is reported for each baseline method.

### Head-to-head SP score comparison with all scalable baselines on QuanTest2 dataset

**Fig. S7.**
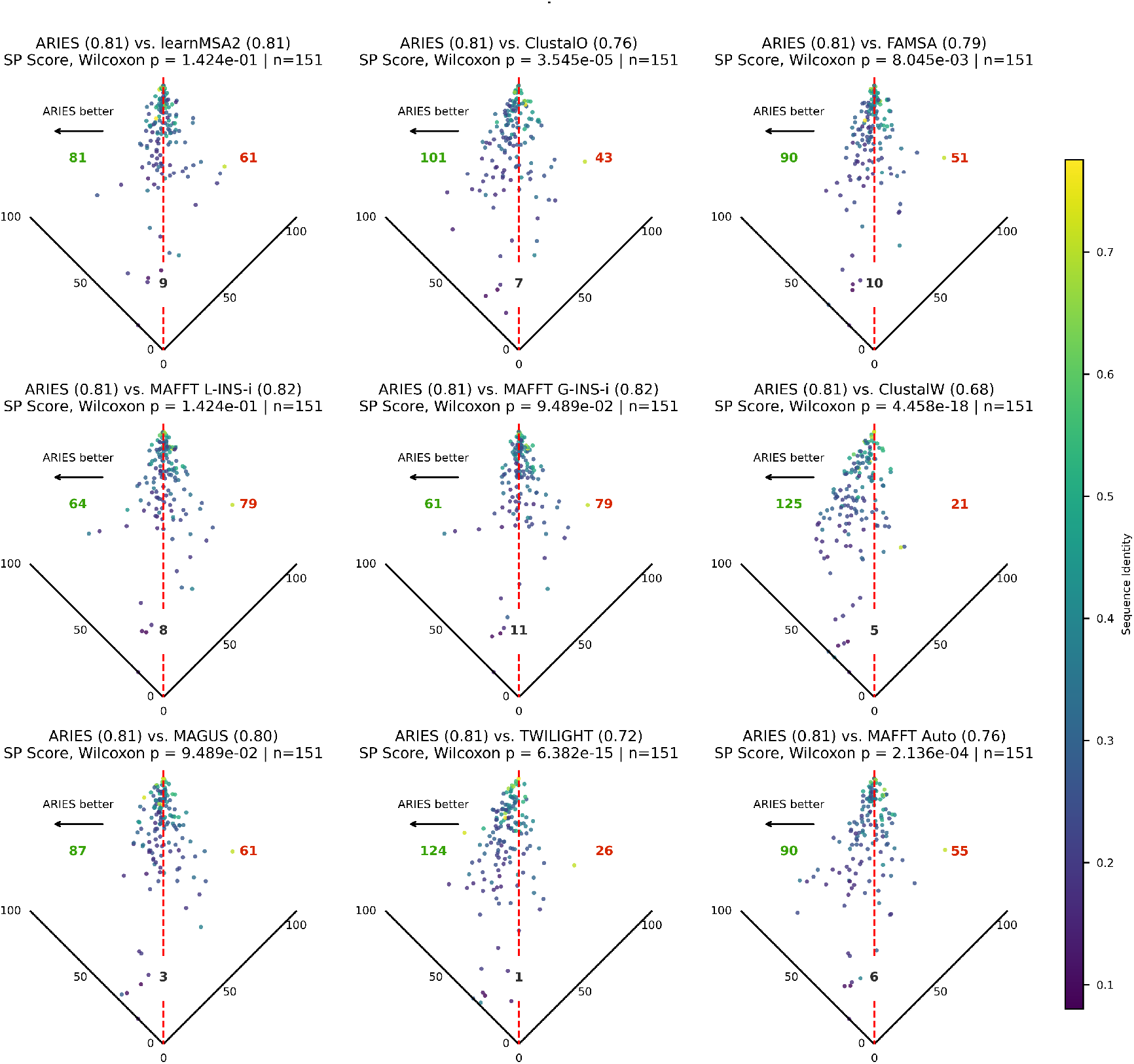
Comparison of ARIES with state-of-the-art MSA methods on QuanTest2 using SP scores; Holm-Bonferroni-corrected *p*-value of one-sided Wilcoxon signed-rank test conducted in the direction of the method with the higher mean SP score is reported for each baseline method.

### Head-to-head TC score comparison with all scalable baselines on QuanTest2 dataset

**Fig. S8.**
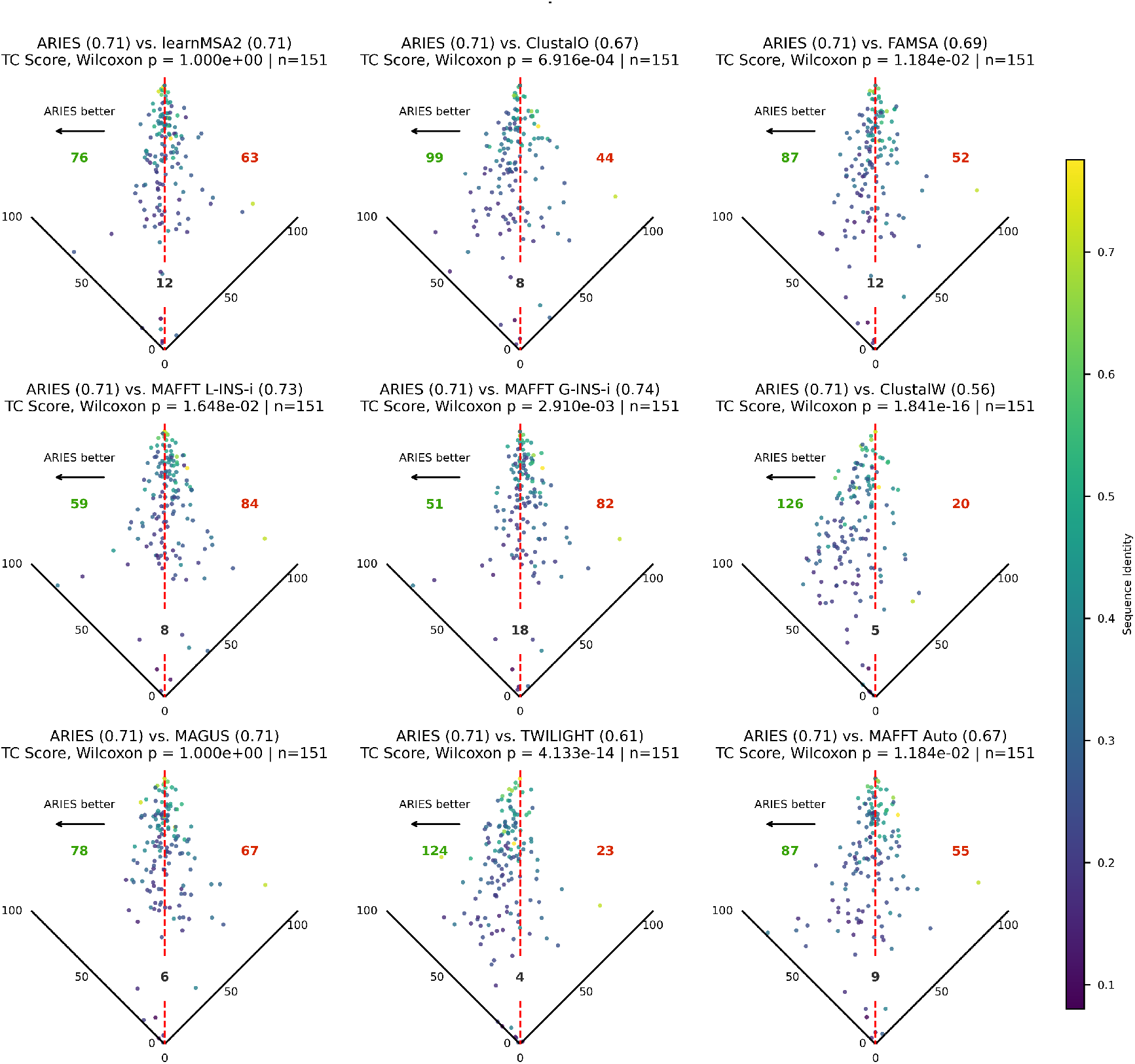
Comparison of ARIES with state-of-the-art MSA methods on QuanTest2 using TC scores; Holm-Bonferroni-corrected *p*-value of one-sided Wilcoxon signed-rank test conducted in the direction of the method with the higher mean TC score is reported for each baseline method.

### SP and TC score comparisons with all baselines on each BAliBASE subset

Tables S2 and S3 summarize comparisons between ARIES and previous methods on each individual BAliBASE reference set (RV11-RV50). Entries denote the mean SP or TC score for each method within a given RV. The sample size (*n*) shown in each column header indicates the number of alignment sets included in that RV. Cells marked with ^*†*^ indicate that one or more alignments failed for that method in the corresponding RV (for example, some methods failed in instances where a sequence contains a non-standard character). In such cases, the comparison with ARIES is performed using only the subset of alignments for which both methods produced valid results. ARIES did not have any failure cases.

Across reference sets, ARIES is strongest on RV11 where it attains higher mean SP and TC scores than every comparison method and wins on a large majority of alignments. ARIES also performs well on RV20, achieving higher mean TC than all comparison methods, and competitive SP performance. On RV12, all methods aside from learnMSA2 achieve uniformly high SP and TC scores, indicating a high-accuracy regime in which performance differences are small in absolute magnitude. Although several refinement-based baselines slightly exceed ARIES in mean score, these differences occur near the performance ceiling and do not reflect large practical gaps. In contrast, RV30 appears more challenging for ARIES: several iterative refinement methods (notably MAFFT variants and MUSCLE) achieve higher mean SP and TC scores and win a larger fraction of alignments. RV40 shows a more balanced pattern, with small mean differences and many non-significant comparisons, suggesting near-parity among methods in that regime. Overall, ARIES demonstrates strong gains in the most divergent setting (RV11), competitive performance in high-accuracy regimes (RV12, RV40), and identifiable opportunities for improvement in insertion-heavy scenarios such as RV30.

**Table S2.** BAliBASE: ARIES vs competitors (SP score) across each RV set in BAliBASE. Entries are mean SP scores. Cells marked ^*†*^ indicate that one or more alignments failed for that method in that RV, so the ARIES comparison used only the subset of alignments that succeeded. The highest score per RV is in bold. In the event of a tie, the method with the lowest *p*-value (one-sided Wilcoxon signed-rank test) is chosen.

| **Method** | **RV11<br>(n=38)** | **RV12<br>(n=44)** | **RV20<br>(n=41)** | **RV30<br>(n=30)** | **RV40<br>(n=49)** | **RV50<br>(n=16)** |
|---|---|---|---|---|---|---|
| ARIES | **0.790** | 0.929 | 0.917 | 0.839 | 0.900 | 0.905 |
| ClustalO | 0.590 | 0.906 | 0.902 | 0.862 | 0.902 | 0.862 |
| ClustalW | 0.501 | 0.864 | 0.852 | 0.725 | 0.789 | 0.743 |
| FAMSA | 0.620 | 0.922 | 0.893 | 0.822 | 0.874 | 0.843 |
| MAFFT G-INS-i | 0.647 | 0.938 | 0.926 | **0.868** | 0.919 | 0.904 |
| MAFFT L-INS-i | 0.651 | 0.937 | **0.927** | 0.860 | **0.919** | 0.902 |
| MAFFT auto | 0.651 | 0.937 | 0.927 | 0.860 | 0.918<sup>†</sup> | 0.902 |
| MAGUS | 0.651 | 0.937 | 0.920 | 0.847 | 0.915 | 0.902 |
| MUSCLE | 0.685 | 0.942 | 0.912 | 0.855 | 0.910 | **0.911** |
| ProbCons | 0.670 | 0.941 | 0.917 | 0.845 | 0.904<sup>†</sup> | 0.894 |
| T-Coffee | 0.658 | **0.944** | 0.915 | 0.838 | 0.906 | 0.895 |
| TWILIGHT | 0.504 | 0.878 | 0.855 | 0.803 | 0.832 | 0.818 |
| learnMSA2 | 0.152 | 0.588 | 0.836<sup>†</sup> | 0.653<sup>†</sup> | 0.656<sup>†</sup> | 0.535 |
| vcMSA | 0.745<sup>†</sup> | 0.920 | 0.857 | 0.798 | 0.733<sup>†</sup> | 0.822 |

**Table S3.** BAliBASE: ARIES vs competitors (TC score) across each RV set in BAliBASE. Entries are mean TC scores. Cells marked ^*†*^ indicate that one or more alignments failed for that method in that RV, so the ARIES comparison used only the subset of alignments that succeeded. The highest score per RV is in bold. In the event of a tie, the method with the lowest *p*-value (one-sided Wilcoxon signed-rank test) is chosen.

| **Method** | **RV11**<br>**(n=38)** | **RV12**<br>**(n=44)** | **RV20**<br>**(n=41)** | **RV30**<br>**(n=30)** | **RV40**<br>**(n=49)** | **RV50**<br>**(n=16)** |
|---|---|---|---|---|---|---|
| ARIES | **0.602** | 0.832 | **0.546** | 0.516 | 0.562 | 0.577 |
| ClustalO | 0.362 | 0.794 | 0.453 | 0.579 | 0.583 | 0.537 |
| ClustalW | 0.230 | 0.717 | 0.222 | 0.276 | 0.398 | 0.312 |
| FAMSA | 0.374 | 0.824 | 0.343 | 0.487 | 0.524 | 0.479 |
| MAFFT G-INS-i | 0.406 | 0.843 | 0.444 | **0.599** | 0.588 | 0.599 |
| MAFFT L-INS-i | 0.424 | 0.845 | 0.468 | 0.586 | **0.607** | 0.597 |
| MAFFT auto | 0.424 | 0.845 | 0.468 | 0.584 | 0.607<sup>†</sup> | 0.595 |
| MAGUS | 0.415 | 0.846 | 0.454 | 0.570 | 0.593 | 0.596 |
| MUSCLE | 0.434 | **0.862** | 0.424 | 0.580 | 0.558 | **0.627** |
| ProbCons | 0.420 | 0.860 | 0.412 | 0.547 | 0.536<sup>†</sup> | 0.579 |
| T-Coffee | 0.410 | 0.859 | 0.406 | 0.474 | 0.554 | 0.587 |
| TWILIGHT | 0.273 | 0.724 | 0.296 | 0.446 | 0.417 | 0.407 |
| learnMSA2 | 0.010 | 0.287 | 0.222<sup>†</sup> | 0.235<sup>†</sup> | 0.152<sup>†</sup> | 0.095 |
| vcMSA | 0.504<sup>†</sup> | 0.824 | 0.436 | 0.387 | 0.405<sup>†</sup> | 0.442 |

## E Additional ablation studies

### Comparing tiled and untiled BAliBASE alignments

**Fig. S9.**
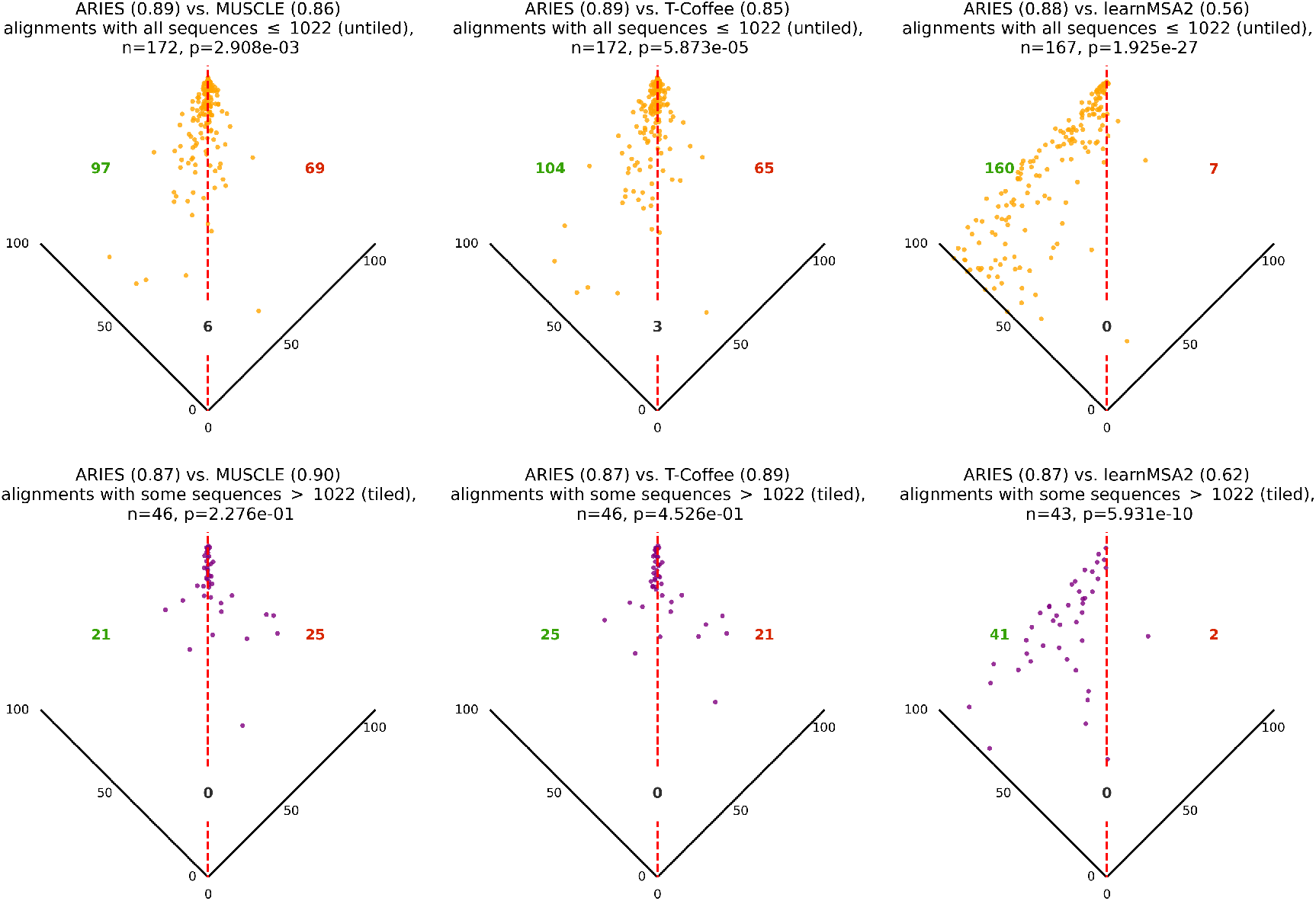
Comparing alignment results for the best-, median-, and lowest-performing competitor methods on BAl-iBASE, for untiled (shown in orange) and tiled (shown in purple) alignments. Any alignment with any sequence longer than 1022 amino acid is considered tiled, regardless of how many sequences in the alignment are actually tiled. We report the Holm–Bonferroni-corrected p-value from a one-sided Wilcoxon signed-rank test, where the tested direction corresponds to the method with the higher mean SP score.

Most existing PLMs are trained with a maximum context length due to the scalability constraints of their training process (Tay et al. 2020). For example, ESM-2 is trained with a maximum context length of 1024 tokens, two of which are reserved for beginning- and end-of-sequence tokens (Rives et al. 2021; Lin et al. 2023). Embeddings for amino acids in longer sequences are typically of lower quality, and running long sequences through PLMs can be prohibitively slow. To work around this constraint, we adopt a tiling strategy that partitions long protein sequences into overlapping segments, thereby preserving contextual continuity across tile boundaries (Brandes et al. 2023). Each tile is processed independently by the PLM, and the resulting representations are subsequently aggregated using a position-weighted average.

This practical trick enables us to align sequence sets with maximum lengths exceeding 1022 residues, covering an additional 46 BAliBASE sets. However, we observe that embeddings derived from tiled sequences tend to be of lower quality than those from untiled sequences, which in turn leads to reduced alignment performance. Figure S9 compares ARIES with the best-, median-, and lowest-performing competitor methods separately on the 172 untiled and 46 tiled BAliBASE sets. While ARIES significantly outperforms all three competitors on the untiled subset, it underperforms the strongest competitor (MUSCLE) on the tiled subset. These findings suggest that ARIES would likely benefit from protein language models trained with inherently longer context windows, which could mitigate the need for tiling.

### Comparing ARIES with vcMSA and learnMSA with the same underlying PLMs

To ensure a fair comparison in which PLM-based baselines employ the same underlying model, we conducted additional experiments comparing ARIES using ProtT5-XL-Half with learnMSA2 (which utilizes ProtT5-XL-Half), as well as ARIES using ProtT5-XL with vcMSA (which utilizes ProtT5-XL) (see Table S4). On the BAliBASE and HOMSTRAD datasets, ARIES significantly out-performs both vcMSA and learnMSA when the same underlying PLM, with all Wilcoxon one-sided signed-rank test *p*-values *<* 1.4 × 10^*−*35^. On QuanTest2, vcMSA was computationally prohibitive and did not complete within a reasonable runtime, and ARIES is slightly outperformed by learn-MSA2, though the difference is not statistically significant (Wilcoxon one-sided signed-rank test, H_1_: learnMSA2 *>* ARIES, yields *p*-value *>* 0.1).

**Table S4.** Across benchmark datasets, we compare the mean SP score of ARIES with Prot-T5-XL to that of vcMSA (which uses Prot-T5-XL), and the mean SP score of ARIES with Prot-T5-XL-Half to that of learnMSA2 (which uses Prot-T5-XL-Half). Within each benchmark dataset, a Holm–Bonferroni correction is applied to correct for multiple comparisons (****, ***, **, and * indicate that the method’s performance differs from ARIES with corrected *p*-values of *≤* 0.0001, 0.001, 0.01, and 0.05, respectively). vcMSA does not scale to QuanTest2 and thus we do not report its mean SP score on that dataset.

| **Baseline (underlying PLM)** | **HOMSTRAD** | **BAliBASE** | **QuanTest2** |
|---|---|---|---|
| ARIES (ProtT5-XL) | **0.90** | **0.86** | **0.78** |
| vcMSA (ProtT5-XL) | 0.87**** | 0.66**** | - |
| ARIES (ProtT5-XL-Half) | **0.90** | **0.86** | 0.78 |
| learnMSA2 (ProtT5-XL-Half) | 0.50**** | 0.48**** | **0.81** |

### Trying different substitution matrices for competitor methods

In the main manuscript, baseline methods were run using their default settings as provided by the authors. For classical protein alignment tools, this corresponds to the BLOSUM62 substitution matrix. For the top three comparison methods by average SP score on each evaluation dataset, we also explored the use of different substitution matrices where supported by the current versions of those tools. These methods included MUSCLE (BAliBASE, HOMSTRAD), MAFFT L-INS-i (BAliBASE, QuanTest2), MAFFT G-INS-i (BAliBASE, QuanTest2), vcMSA (HOMSTRAD), T-Coffee (HOMSTRAD), and learnMSA2 (QuanTest2).

Among these, only MAFFT L-INS-i, MAFFT G-INS-i, and T-Coffee provide support in their current releases for specifying alternative amino acid substitution matrices. Other methods either do not expose this option (e.g., MUSCLE, MAGUS), or rely on fixed internal scoring schemes or downstream alignment components that do not directly admit user-specified matrices in practice. Accordingly, on datasets where one of these matrix-configurable methods ranked among the top three, we evaluated performance under two additional substitution matrices.

For BAliBASE, both MAFFT L-INS-i and MAFFT G-INS-i ranked among the top three and allow substitution matrix changes. We tested BLOSUM45 and JTT250 in addition to the default BLOSUM62. Across matrices, ARIES significantly outperforms both MAFFT G-INS-i and MAFFT L-INS-i, shown in Fig. S10.

For HOMSTRAD, T-Coffee was the only top-three method supporting matrix selection. We evaluated BLOSUM45 and PAM250 (as JTT matrices are not supported in this configuration) in addition to the default BLOSUM62. ARIES significantly outperforms T-Coffee regardless of the substitution matrix used (Wilcoxon signed-rank test corrected p-values all *<* 1*e*^*−*25^), shown in Fig. S11.

For QuanTest2, both MAFFT L-INS-i and MAFFT G-INS-i ranked among the top three and allow substitution matrix changes. We again tested BLOSUM45 and JTT250 in addition to the default BLOSUM62. Across all three matrices, performance differences were small and inconsistent. In some configurations, ARIES produced higher SP scores on more datasets, while in others, the MAFFT variants did. However, none of these differences were statistically significant (corrected one-sided Wilcoxon signed-rank test p-values all *>* 0.1), shown in Fig. S12. Thus, varying the substitution matrix did not materially affect the relative performance of the methods.

**Fig. S10.**
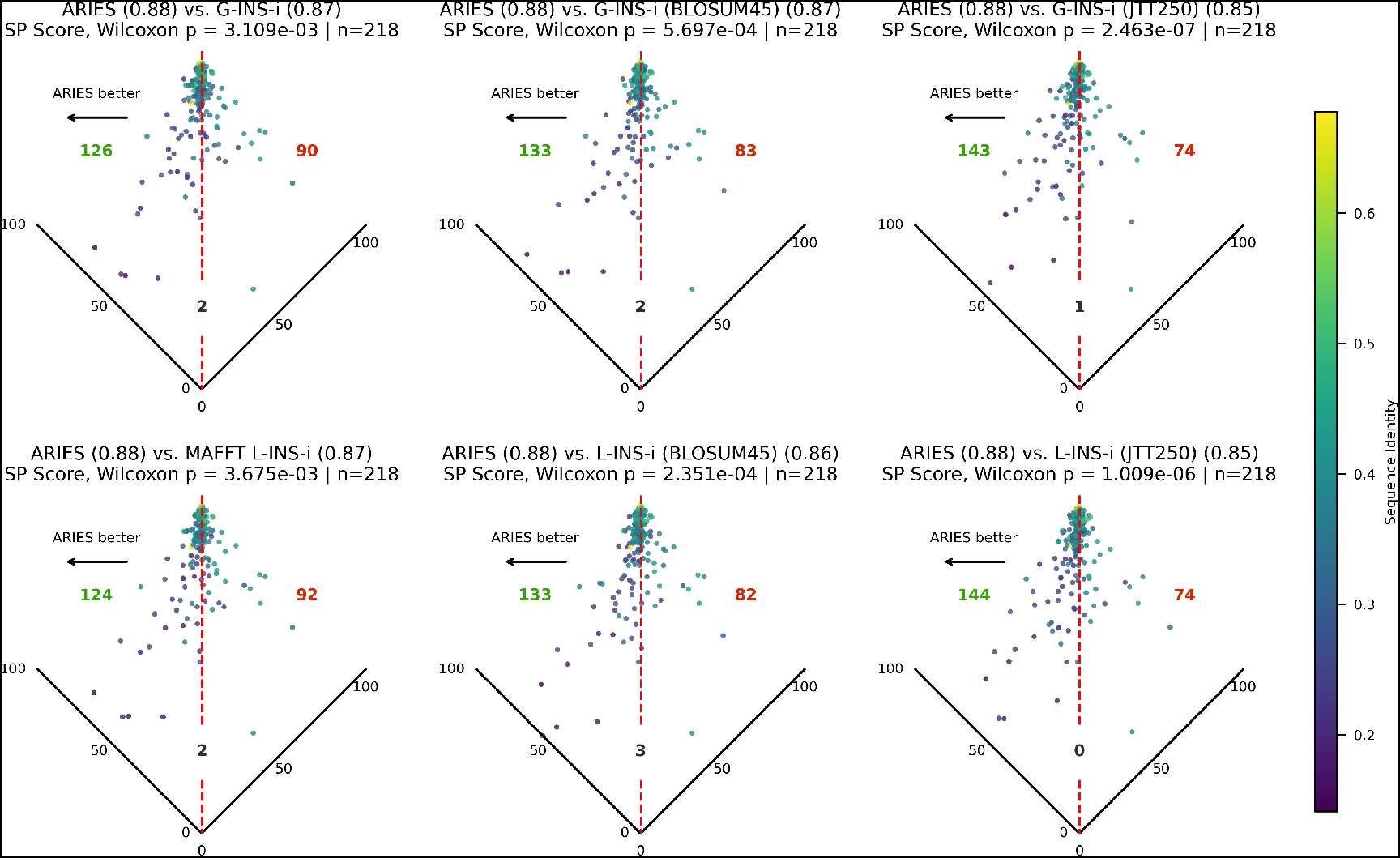
SP scores across all BAliBASE sets of ARIES, MAFFT L-INS-i, and MAFFT G-INS-i with different substitution matrices. ARIES significantly outperforms both MAFFT variants regardless of substitution matrix.

Across all tested configurations, changing substitution matrices did not alter the conclusions of our study.

### Comparing star-alignment ARIES with progressive-alignment ARIES

In our PLM-based progressive alignment framework, one must construct an embedding-level representation for each intermediate aligned cluster before proceeding to the next merge; this is due to the DTW procedure taking as input amino acid embeddings. Thus, this requires generating a new embedding representation at each stage in the progressive alignment. We implemented two progressive alignment variants of ARIES. Specifically, we used the ClustalW guide tree to determine the order of alignment. Initially, each sequence is treated as an individual cluster. At every progressive step, a pair of clusters is aligned via dynamic time warping (DTW) and merged into a new cluster. The embedding of the newly formed cluster is computed in one of two ways: (1) reembedding the aligned protein sequences with the PLM, where gap characters are replaced by ‘X’ tokens, similar to how ARIES synthesizes its template embeddings (we refer to this progressive alignment version as “Re-Embed”), or (2) simply averaging the embeddings of aligned residues, while keeping insertion embeddings unchanged from their original sequence (we refer to this version as “Mean-Merge”). When aligning clusters, gaps are handled conservatively: once introduced, a gap is preserved according to its earliest occurrence.

**Fig. S11.**
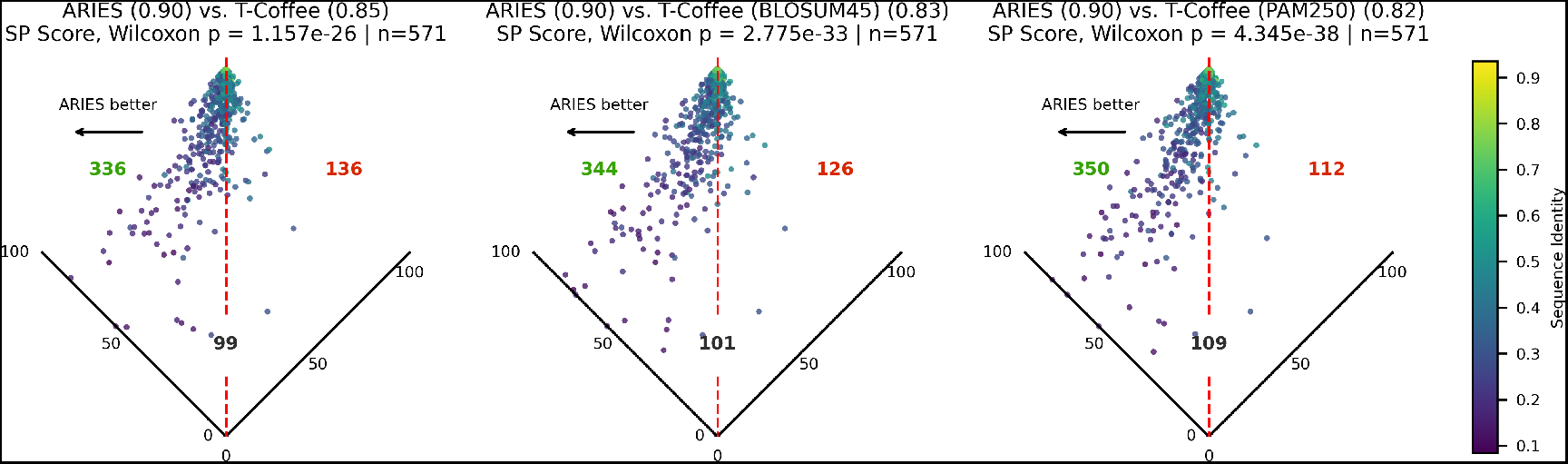
SP scores across all HOMSTRAD sets of ARIES and T-Coffee with different substitution matrices. ARIES significantly outperforms T-Coffee in all tested configurations.

**Fig. S12.**
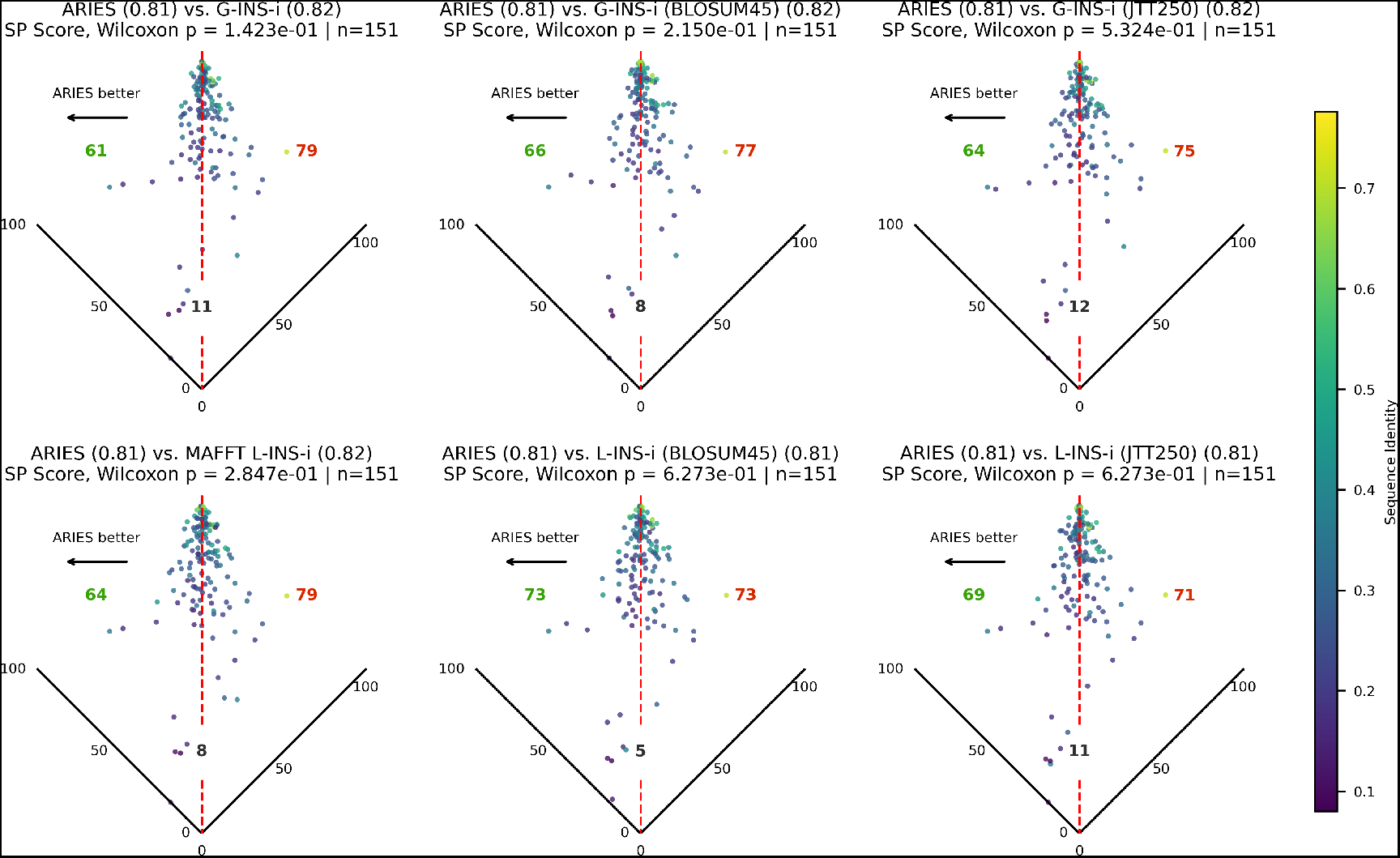
SP scores across all QuanTest2 sets of ARIES, MAFFT L-INS-i, and MAFFT G-INS-i with different substitution matrices. *p*-values shown correspond to one-sided Wilcoxon signed-rank tests for the method with the higher average SP score outperforming the other, with a Holm-Bonferroni correction.

Fig. S13 presents volcano plots comparing the SP scores achieved by star-alignment ARIES compared to these variants on all BAliBASE alignment sets. ARIES with star-alignment achieves an average SP score of 0.72, significantly outperforming both the Re-Embed progressive alignment (average SP score 0.68, *p* = 5.20*e*^*−*10^) and the Mean-Merge progressive alignment (average SP score 0.69, *p* = 1.32*e*^*−*2^). We stress that we do not claim that progressive alignment strategies are inherently inferior. Alternate approaches for cluster embedding generation, or incorporating strategies such as iterative refinement, may improve progressive alignment performance and would be a promising direction for future work.

**Fig. S13.**
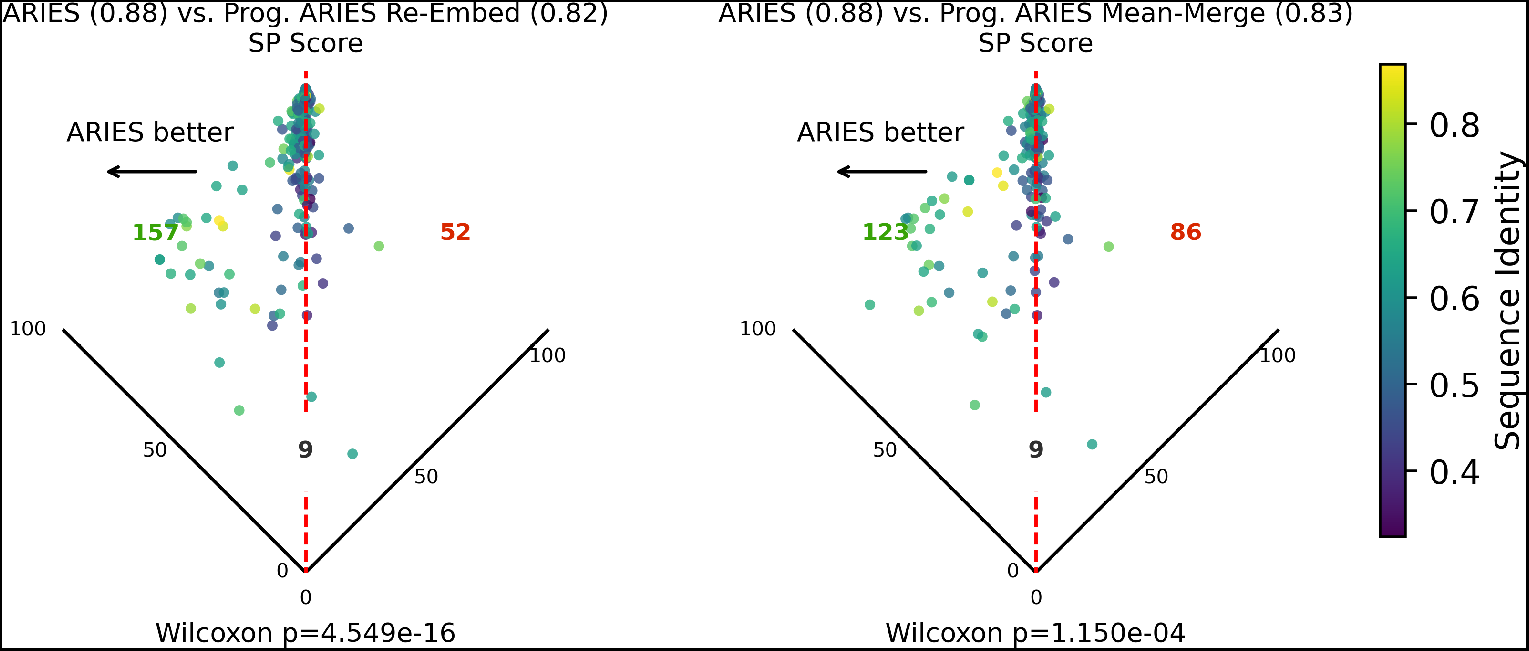
Average SP score (across all BAliBASE sets) of ARIES with star-vs. two progressive alignment strategies.

### On the optimal number of medoids for ARIES template synthesis

To assess whether *K* = ⌈ln(*N* )⌉ serves as an effective heuristic that frequently yields near-optimal performance, we reproduce Fig. 5 for all 151 QuanTest MSA sets, subsampling each alignment to the first *N* = 250, 500, 750, 1000 sequences. The y-axis is normalized against the mean SP score of the best *K* to compare across different values of *N* . Although *K* = ⌈ln(*N* )⌉ (highlighted in red) does not always attain the absolute best performance, it is consistently within 1.6% of the best performance for *K* ∈ {1, 2, …, 20} across all values of *N* .

**Fig. S14.**
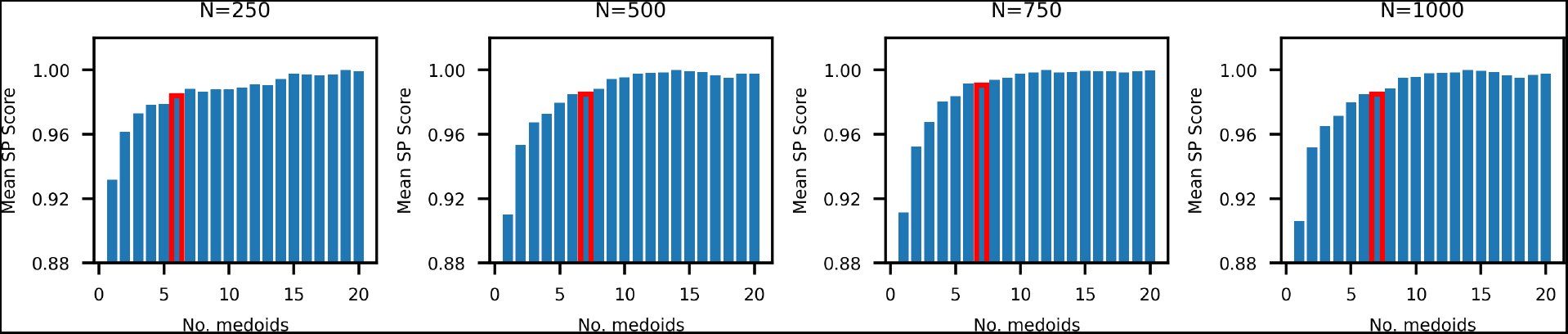
Normalized average SP score (across all BAliBASE sets) of ARIES with different no. top-*K* medoids (*K ∈ {*1, 2, …, 20*}*) for template construction. *K* = *⌈*ln(*N* )*⌉* is higlighted in red.

### Runtime comparison at different input sizes

**Fig. S15.**
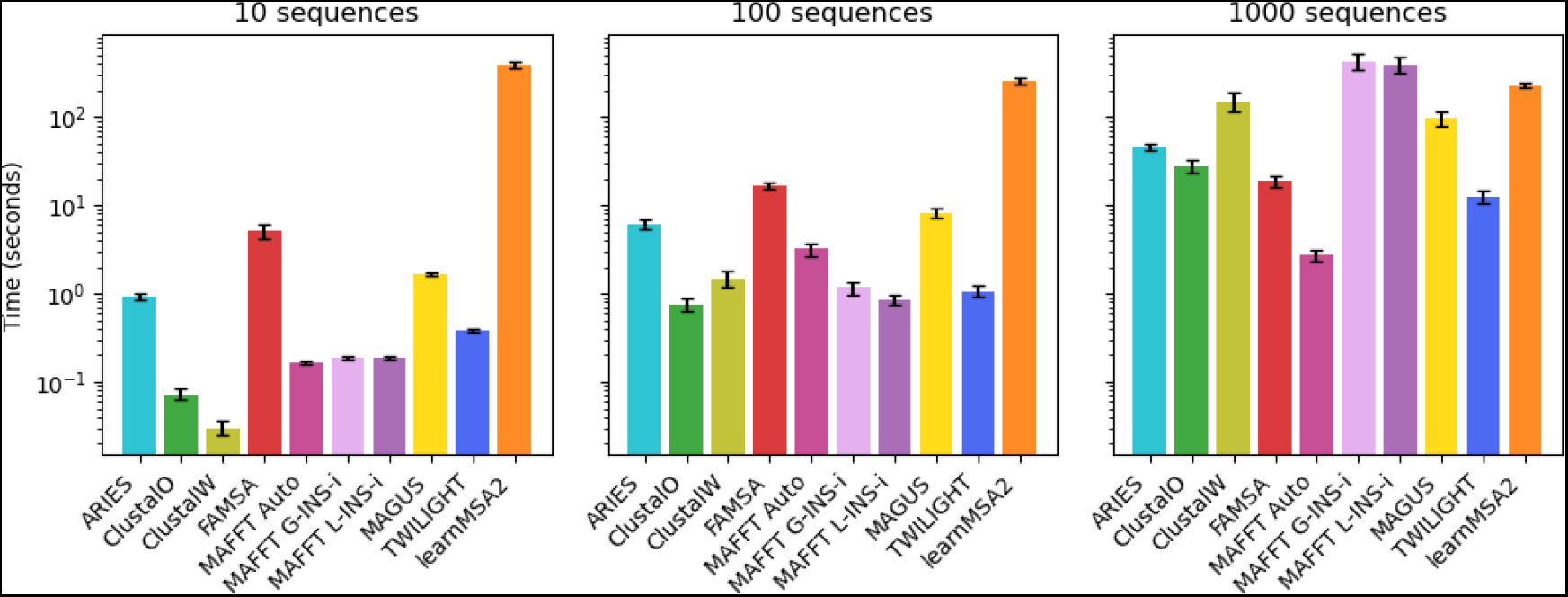
Runtime comparison of 10 MSA methods on QuanTest2 sets of 10, 100, and 1000 sequences.

Among the methods able to produce alignments of 1000 sequences in under 10 minutes on average, we measured runtime on sequence sets of size 10, 100, and 1000 (Fig. S15). These sets are the first *N* sequences in each QuanTest2 reference alignment set. We note again that each baseline was tested using the default arguments to maintain consistency. Some tools (e.g., MUSCLE) have faster modes, but we did not enable these non-default modes to ensure fair comparison across tools. All CPU-based baseline methods were run with 10 CPU cores and 10 GB of memory percore. GPU-accelerated methods (ARIES and learnMSA2) were executed on a single NVIDIA A100 GPU with 80 GB of memory and the same CPU limits. For ARIES, we select the ESM-2 model with 650M parameters with embedding depth 9 based on our ablation results, despite the higher runtime compared to smaller or shallower configurations. Importantly, even with this more expensive setting, our method remains substantially faster than the other GPU-based competitor, learnMSA2.

ARIES scales approximately linearly with *N*, as do several of the CPU-based baselines. learnMSA2, which is the only other GPU-enabled method we evaluate, has a runtime that scales linearly with *N* but requires roughly the same runtime on small sequence sets as it does on sets of 1000 sequences likely because it struggles to converge when only a few samples are available.

### Runtime comparison using ARIES for varying embedding depths and dimensions

Fig. S16 examines the impact of model size and embedding depth on the runtime of ARIES. We compare two ESM variants, with 150M and 650M parameters, respectively. The results indicate that, within the same class of underlying PLM, ARIES runtime scales linearly with both the number of model parameters and the embedding depth.

**Fig. S16.**
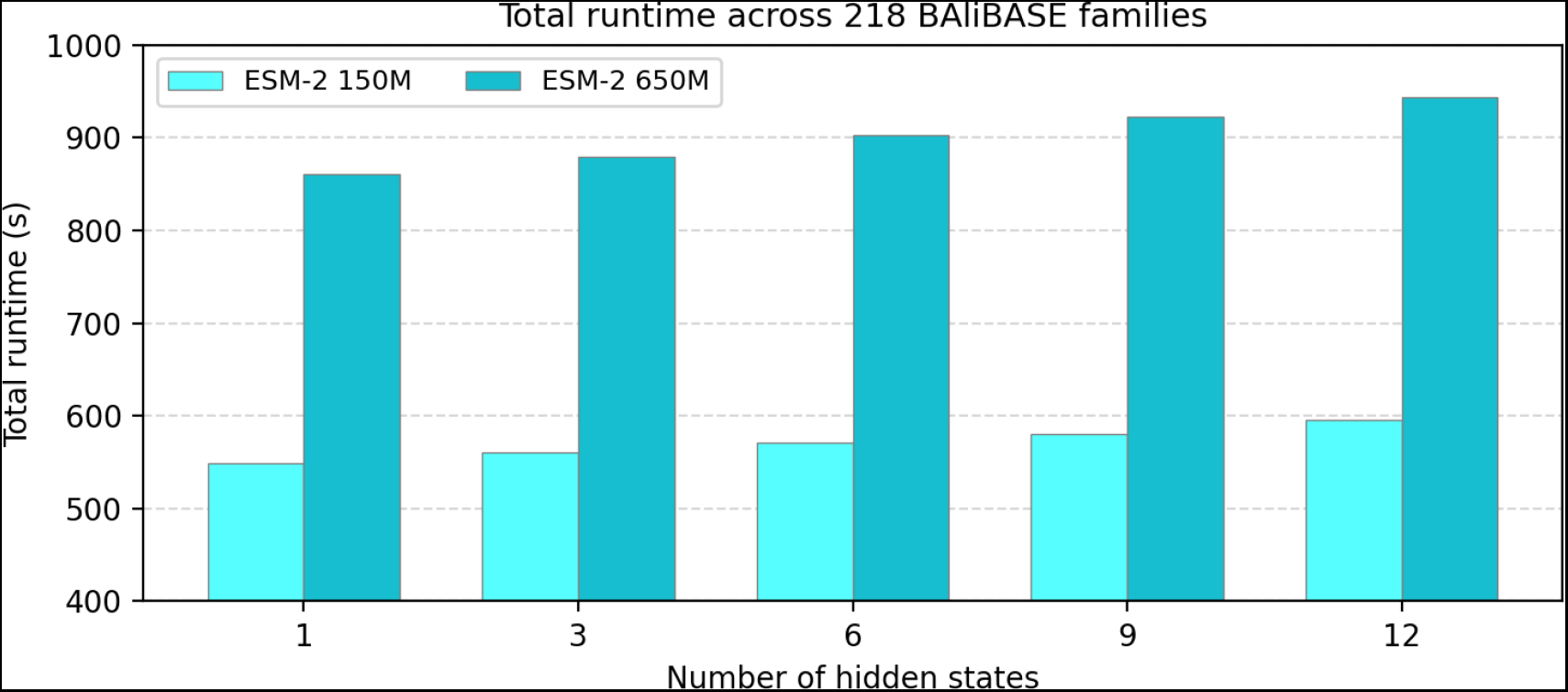
Runtime comparison for the entire BAliBASE dataset. Larger PLMs are substantially slower, and runtime increases further with embedding depth, though to a lesser extent.

## Bibliography

Alley E, Khimulya G, Biswas S, AlQuraishi M, Church G. 2019. Unified rational protein engineering with sequence-based deep representation learning. Nature Methods 16: 1315–1322.

Altschul S, Lipman D. 1989. Trees, stars, and multiple biological sequence alignment. SIAM Journal of Applied Mathematics 49: 197–209.

Altschul SF. 1991. Amino acid substitution matrices from an information theoretic perspective. Journal of Molecular Biology 219(3): 555–565. ISSN 0022-2836.

Altschul SF, Gish W, Miller W, Myers EW, Lipman DJ. 1990. Basic local alignment search tool. Journal of Molecular Biology 215(3): 403–410.

Becker F, Stanke M. 2024. learnMSA2: deep protein multiple alignments with large language and hidden Markov models. Bioinformatics 40(Supplement 2): ii79–ii86. ISSN 1367-4811.

Bepler T, Berger B. 2021. Learning the protein language: Evolution, structure, and function. Cell Systems 12(6): 654–669.

Bepler T, Berger B. 2019. Learning protein sequence embeddings using information from structure. Proceedings of International Conference on Learning Representations. URL http://arxiv.org/abs/1902.08661.

Biegert A, Söding J. 2009. Sequence context-specific profiles for homology searching. Proceedings of the National Academy of Sciences 106(10): 3770–3775. doi: 10.1073/pnas.0810767106.

Brandes N, Goldman G, Wang CH, Ye CJ, Ntranos V. 2023. Genome-wide prediction of disease variant effects with a deep protein language model. Nature Genetics 55(9): 1512–1522.

Capra JA, Singh M. 2007. Predicting functionally important residues from sequence conservation. Bioinformatics 23(15): 1875–1882.

Dayhoff M, Schwartz R, Orcutt B. 1978. A model of evolutionary change in proteins. Atlas of protein sequence and structure 5: 345–352.

Deorowicz S, Debudaj-Grabysz A, Gudyś A. 2016. FAMSA: Fast and accurate multiple sequence alignment of huge protein families. Scientific Reports 6(1): 33964. ISSN 2045-2322.

Do CB, Mahabhashyam MS, Brudno M, Batzoglou S. 2005. ProbCons: Probabilistic consistency-based multiple sequence alignment. Genome Research 15(2): 330–340. ISSN 1088-9051.

Edgar RC. 2004. MUSCLE: a multiple sequence alignment method with reduced time and space complexity. BMC Bioinformatics 5(1): 113. ISSN 1471-2105.

Elnaggar A, Heinzinger M, Dallago C, Rehawi G, Yu W, Jones L, Gibbs T, Feher T, Angerer C, Steinegger M, et al. 2021. ProtTrans: Towards cracking the language of life’s code through self-supervised learning. IEEE Transactions on Pattern Analysis and Machine Intelligence.

Henikoff S, Henikoff JG. 1992. Amino acid substitution matrices from protein blocks. Proceedings of the National Academy of Sciences of the United States of America 89(22): 10915–10919. ISSN 0027-8424.

Hoang M, Singh M. 2025. Locality-aware pooling enhances protein language model performance across varied applications. Bioinformatics 41(Supplement 1): i217–i226.

Hoang M, Zheng H, Kingsford C. 2022. Deepminimizer: A differentiable framework for optimizing sequence-specific minimizer schemes. In International Conference on Research in Computational Molecular Biology, 52–69. Springer.

Hoang M, Marçais G, Kingsford C. 2024. Density and conservation optimization of the generalized masked-minimizer sketching scheme. Journal of Computational Biology 31(1): 2–20.

Iovino BG, Ye Y. 2024. Protein embedding based alignment. BMC Bioinformatics 25(1): 85. ISSN 1471-2105.

Iovino BG, Tang H, Ye Y. 2024. Protein domain embeddings for fast and accurate similarity search. Genome Research 34(9): 1434–1444. ISSN 1088-9051, 1549-5469.

Jumper J, Evans R, Pritzel A, Green T, Figurnov M, Ronneberger O, Tunyasuvunakool K, Bates R, Žídek A, Potapenko A, et al. 2021. Highly accurate protein structure prediction with AlphaFold. Nature 596(7873): 583–589. ISSN 1476-4687.

Kaminski K, Ludwiczak J, Pawlicki K, Alva V, Dunin-Horkawicz S. 2023. pLM-BLAST: distant homology detection based on direct comparison of sequence representations from protein language models. Bioinformatics 39(10): btad579. ISSN 1367-4811.

Katoh K, Standley DM. 2013. MAFFT Multiple Sequence Alignment Software Version 7: Improvements in Performance and Usability. Molecular Biology and Evolution 30(4): 772–780. ISSN 0737-4038.

Katoh K, Misawa K, Kuma K, Miyata T. 2002. MAFFT: a novel method for rapid multiple sequence alignment based on fast Fourier transform. Nucleic Acids Research 30(14): 3059–3066. ISSN 0305-1048.

Kilinc M, Jia K, Jernigan RL. 2023. Improved global protein homolog detection with major gains in function identification. Proceedings of the National Academy of Sciences 120(9): e2211823120.

Krogh A, Brown M, Mian IS, Sjölander K, Haussler D. 1994. Hidden markov models in computational biology: Applications to protein modeling. Journal of Molecular Biology 235(5): 1501–1531.

Lin Z, Akin H, Rao R, Hie B, Zhu Z, Lu W, Smetanin N, Verkuil R, Kabeli O, Shmueli Y, et al. 2023. Evolutionary-scale prediction of atomic-level protein structure with a language model. Science 379(6637): 1123–1130.

Liu K, Warnow TJ, Holder MT, Nelesen SM, Yu J, Stamatakis AP, Linder CR. 2012. SATe-II: very fast and accurate simultaneous estimation of multiple sequence alignments and phylogenetic trees. Systematic Biology 61(1): 90–106. ISSN 1076-836X.

Llinares-López F, Berthet Q, Blondel M, Teboul O, Vert JP. 2023. Deep embedding and alignment of protein sequences. Nature Methods 20(1): 104–111.

Lupo U, Sgarbossa D, Bitbol AF. 2022. Protein language models trained on multiple sequence alignments learn phylogenetic relationships. Nature Communications 13(1): 6298.

Marçais G, Pellow D, Bork D, Orenstein Y, Shamir R, Kingsford C. 2017. Improving the performance of minimizers and winnowing schemes. Bioinformatics 33(14): i110–i117.

Marks DS, Hopf TA, Sander C. 2012. Protein structure prediction from sequence variation. Nature Biotechnology 30(11): 1072–1080.

McWhite CD, Armour-Garb I, Singh M. 2023. Leveraging protein language models for accurate multiple sequence alignments. Genome Research 33(7): 1145–1153. ISSN 1088-9051.

Mirarab S, Nguyen N, Guo S, Wang LS, Kim J, Warnow T. 2015. PASTA: Ultra-Large Multiple Sequence Alignment for Nucleotide and Amino-Acid Sequences. Journal of Computational Biology 22(5): 377–386. ISSN 1557-8666.

Mistry J, Chuguransky S, Williams L, Qureshi M, Salazar G, Sonnhammer E, Tosatto S, Paladin L, Raj S, Richardson L, et al. 2021. Pfam: The protein families database in 2021. Nucleic Acids Research 49: D412–D419.

Needleman SB, Wunsch CD. 1970. A general method applicable to the search for similarities in the amino acid sequence of two proteins. Journal of Molecular Biology 48(3): 443–453. ISSN 0022-2836.

Notredame C, Higgins DG, Heringa J. 2000. T-coffee: a novel method for fast and accurate multiple sequence alignment1. Journal of Molecular Biology 302(1): 205–217. ISSN 0022-2836.

Pantolini L, Studer G, Pereira J, Durairaj J, Tauriello G, Schwede T. 2024. Embedding-based alignment: combining protein language models with dynamic programming alignment to detect structural similarities in the twilight-zone. Bioinformatics 40(1): btad786. ISSN 1367-4811.

Petti S, Bhattacharya N, Rao R, Dauparas J, Thomas N, Zhou J, Rush AM, Koo P, Ovchinnikov S. 2023. End-to-end learning of multiple sequence alignments with differentiable Smith–Waterman. Bioinformatics 39(1): btac724.

Rao R, Bhattacharya N, Thomas N, Duan Y, Chen P, Canny J, et al. 2019. Evaluating protein transfer learning with TAPE. In Advances in Neural Information Processing Systems (ed. H Wallach, H Larochelle, A Beygelzimer, F d’Alché-Buc, E Fox, R Garnett), volume 32.

Remmert M, Biegert A, Hauser A,, Söding J. 2012. HHblits: lightning-fast iterative protein sequence searching by HMM-HMM alignment. Nature Methods 9: 73–175.

Rives A, Meier J, Sercu T, Goyal S, Lin Z, Liu J, Guo D, Ott M, Zitnick CL, Ma J, et al. 2021. Biological structure and function emerge from scaling unsupervised learning to 250 million protein sequences. Proceedings of the National Academy of Sciences 118(15): e2016239118.

Rost B. 1999. Twilight zone of protein sequence alignments. Protein Engineering 12(2): 85–94. ISSN 0269-2139.

Sakoe H, Chiba S. 1978. Dynamic programming algorithm optimization for spoken word recognition. IEEE Transactions on Acoustics, Speech, and Signal Processing 26(1): 43–49. ISSN 0096-3518.

Schütze K, Heinzinger M, Steinegger M, Rost B. 2022. Nearest neighbor search on embeddings rapidly identifies distant protein relations. Frontiers in Bioinformatics 2. ISSN 2673-7647.

Sievers F, Higgins DG. 2019. Quantest2: benchmarking multiple sequence alignments using secondary structure prediction. Bioinformatics 36(1): 90–95. ISSN 1367-4803.

Sievers F, Wilm A, Dineen D, Gibson TJ, Karplus K, Li W, Lopez R, McWilliam H, Remmert M, Söding J, et al. 2011. Fast, scalable generation of high-quality protein multiple sequence alignments using Clustal Omega. Molecular Systems Biology 7: 539. ISSN 1744-4292.

Sleator DD, Tarjan RE. 1981. A data structure for dynamic trees. In Proceedings of the Thirteenth Annual ACM Symposium on Theory of Computing, 114–122.

Smirnov V, Warnow T. 2021. MAGUS: multiple sequence alignment using graph clustering. Bioinformatics 37(12): 1666–1672.

Sokal RR, Michener CD, et al. 1958. A statistical method for evaluating systematic relationships. University of Kansas Scientific Bulletin 38: 1409–1438.

Stebbings LA, Mizuguchi K. 2004. Homstrad: recent developments of the homologous protein structure alignment database. Nucleic Acids Research 32: D203–D207. ISSN 0305-1048.

Tay Y, Dehghani M, Abnar S, Shen Y, Bahri D, Pham P, Rao J, Yang L, Ruder S, Metzler D. Long range arena: A benchmark for efficient transformers, 2020. URL https://arxiv.org/abs/2011.04006.

Thompson JD, Higgins DG, Gibson TJ. 1994. CLUSTAL W: improving the sensitivity of progressive multiple sequence alignment through sequence weighting, position-specific gap penalties and weight matrix choice. Nucleic Acids Research 22(22): 4673–4680. ISSN 0305-1048.

Thompson JD, Koehl P, Ripp R, Poch O. 2005. Balibase 3.0: Latest developments of the multiple sequence alignment benchmark. Proteins. 61(1). ISSN 0887-3585.

Tseng YH, Walia S, Turakhia Y. 2025. Ultrafast and ultralarge multiple sequence alignments using twilight. Bioinformatics 41(Supplement 1): i332–i341.

Whelan S, Goldman N. 2001. A general empirical model of protein evolution derived from multiple protein families using a maximum-likelihood approach. Molecular Biology and Evolution 18(5): 691–699.

